# The Biomechanical Basis of Biased Epithelial Tube Elongation in Lung and Kidney Development

**DOI:** 10.1101/2020.06.22.166231

**Authors:** Lisa Conrad, Steve Runser, Harold Gómez, Christine Lang, Mathilde Dumond, Aleksandra Sapala, Laura Kramps, Odysse Michos, Roman Vetter, Dagmar Iber

## Abstract

During lung development, epithelial branches expand preferentially in longitudinal direction. This bias in outgrowth has been linked to a bias in cell shape and in the cell division plane. How this bias arises is unknown. Here, we show that biased epithelial outgrowth occurs independent of the surrounding mesenchyme, of preferential turn-over of the extracellular matrix at the bud tips, and of FGF signalling. There is also no evidence for actin-rich filopodia at the bud tips. Rather, we find epithelial tubes to be collapsed during early lung and kidney development, and we observe fluid flow in the narrow tubes. By simulating the measured fluid flow inside segmented narrow epithelial tubes, we show that the shear stress levels on the apical surface are sufficient to explain the reported bias in cell shape and outgrowth. We use a cell-based vertex model to confirm that apical shear forces, unlike constricting forces, can give rise to both the observed bias in cell shapes and tube elongation. We conclude that shear stress may be a more general driver of biased tube elongation beyond its established role in angiogenesis.

**Summary Statement:** We systematically analysed the requirements for biased elongating outgrowth of epithelial tubes during lung and kidney development, and provide evidence that fluid-flow induced shear stress drives their biased elongation.

## INTRODUCTION

Epithelial tubes are an essential component of many organs. During development, epithelial tubes elongate (Fig. 1A). Tube elongation can be either isotropic or anisotropic, i.e. the tubes either lengthen as much as they widen, or there is a bias in outgrowth (Fig. 1B). Growth is by default isotropic, and a bias in elongation can therefore only arise if growth symmetry is broken in the epithelium. How this symmetry break is achieved is largely elusive. We focus here on the mouse embryonic lung and kidney. In the mouse lung, epithelial tube expansion is anisotropic initially (E10.5-E11.5), but, at least in the trachea, becomes isotropic at later stages (from E12.5) (Kishimoto et al., 2018; Tang et al., 2011; Tang et al., 2018). The biased outgrowth has been related to a bias in the orientation of the mitotic spindles of dividing cells (Saburi et al., 2008; Tang et al., 2011; Tang et al., 2018; Yates et al., 2010). According to Hertwig’s rule (Hertwig, 1884), cells divide through their mass point and perpendicular to their longest axis. Indeed, the bias in cell division is accompanied by a bias in cell shape (Tang et al., 2011; Tang et al., 2018). The planar cell polarity (PCP) pathway plays an important role in regulating the mitotic spindle angle distribution in many organs, including the embryonic renal tubes (Ciruna et al., 2006; Gong et al., 2004; Saburi et al., 2008), though no such involvement could be ascertained for the early stages of lung development (Tang et al., 2011). Independent of whether the PCP pathway is involved, it remains an open question how the elongation bias and its direction arise in the first place.

**Figure 1.**
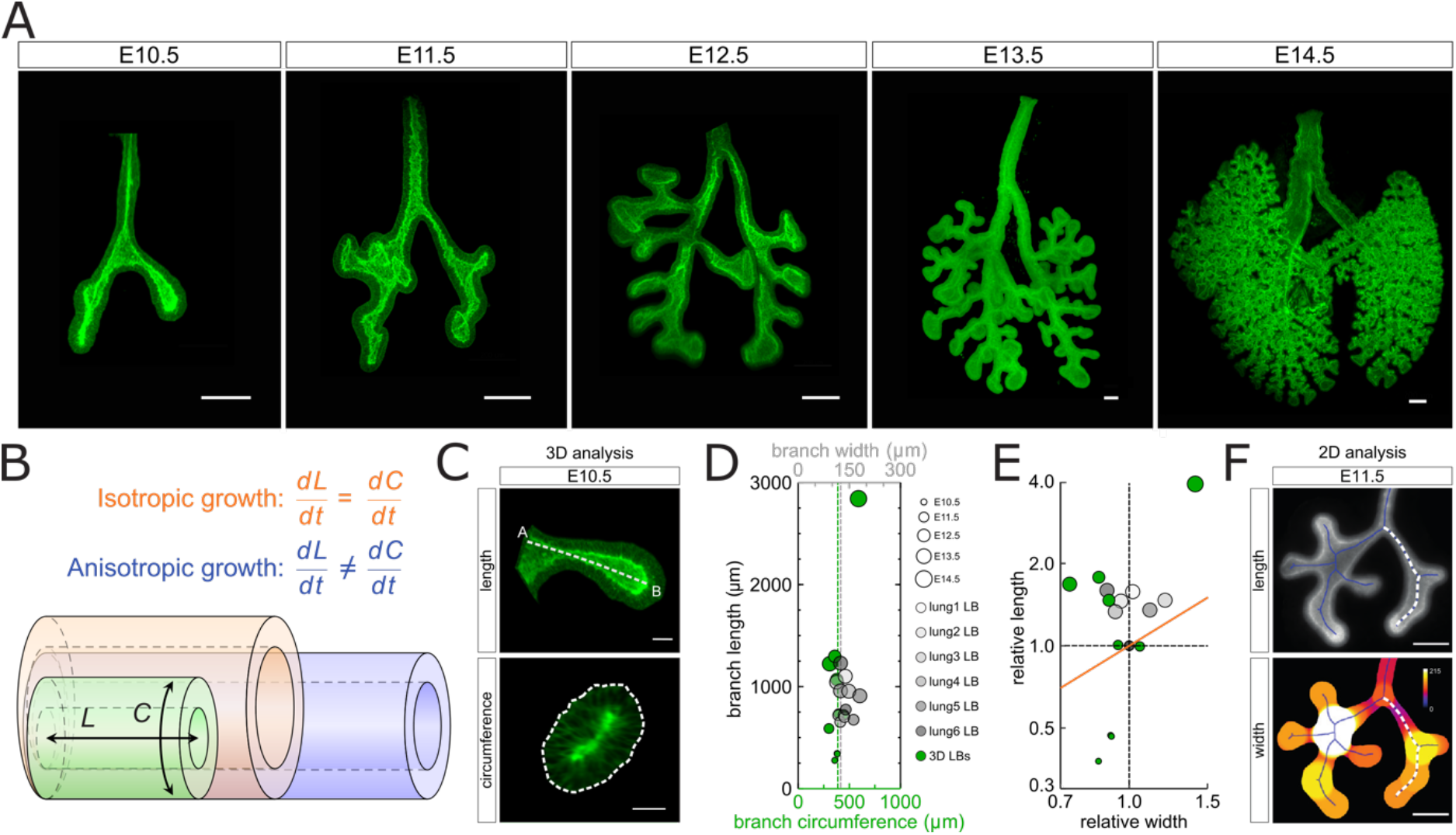
Biased epithelial lung tube elongation. **(A)** Developmental timeline of serial dissections from mouse embryonic lungs expressing the Shh^GC/+^; ROSA^mT/mG^ reporter (green epithelium, and red mesenchyme) imaged using light-sheet microscopy. Scale bars 200 μm. **(B)** Schematic of isotropic and anisotropic tube expansion during development. **(C)**3D morphometric measurements of epithelial tube length and circumference for an E10.5 left bronchus. Scale bars 50 μm. **(D)** 3D length and average circumference measurements of the left bronchus of embryonic lungs (E10.5-E14.5 in green), and 2D length and diameter measurements for E11.5 lungs cultured on a filter over 48h (grey). Dot size increases with the developmental stage. **(E)** Relative width and length for 3D serial dissections (green) and 2D filter-cultured lungs (grey), normalised to their average size at E11.5. Between E10.5 and E11.5 there is roughly 2-fold more lengthening than widening. After E11.5, data points remain above the orange line, implying that branches continue to elongate more than they widen (anisotropic elongation). Between E13.5 and E14.5, expansion is roughly isotropic. **(F)** 2D morphometric measurements of length and diameter for a filter-cultured E11.5 lung. Width scale (colour bar) in μm. Scale bars 200 μm.

In principle, a bias in outgrowth could originate from polarization along the tube, from a pulling force at the tip, or from a mechanical constraint that limits expansion in the circumferential direction. Several signalling pathways are known to affect the bias in lung tube elongation. Thus, hyperactive KRas (KRas^G12D^) in the lung epithelium abrogates the bias in outgrowth during lung branching morphogenesis (Tang et al., 2011), and pharmacological reagents that activate or inhibit fibroblastic growth factor (FGF) signalling, sonic hedgehog (SHH) signalling, or L-type Ca2+ channels affect the width of cultured lung buds (Goodwin et al., 2019). FGF10 and Glial cell line-derived neurotrophic factor (GDNF) signalling are necessary for the formation of branches in the lung and kidney, respectively (Michos et al., 2010; Min et al., 1998; Moore et al., 1996; Pichel et al., 1996; Rozen et al., 2009; Sanchez et al., 1996; Sekine et al., 1999). FGF10 has been proposed to act as chemoattractant because it is secreted from the submesothelial mesenchyme, and isolated lung epithelia grow towards an FGF10 source (Park et al., 1998). However, *Gdnf* is expressed uniformly in the ureteric cap mesenchyme (Hellmich et al., 1996), and branching morphogenesis is still observed when *Fgf10* is expressed uniformly in the lung mesenchyme (Volckaert et al., 2013), which contradicts a need for a chemoattractant gradient. In both cases, a Turing mechanism may lead to the emergence of focused FGF10-FGFR2b and GDNF-RET signalling at the branch points, despite the uniform expression of the ligand (Menshykau et al., 2014; Menshykau et al., 2019). Focused signalling could then, in principle, result in epithelial leader cells as have been identified in the *Drosophila* trachea and the mammalian kidney (Chi et al., 2009; Ghabrial and Krasnow, 2006). Also, the mesenchymal tip could, in principle, drive elongation as observed in the chick Wolffian duct, where FGF8 prevents a transition to an epithelial state and drives tissue elongation by promoting migratory and motile properties in the mesenchymal tip cells (Atsuta and Takahashi, 2015).

Alternatively, mechanical constraints may drive tube elongation, as proposed for the mammary gland (Paine et al., 2016). Lungs and kidneys do not have a myoepithelium like mammary glands, but smooth muscles could, in principle, drive tube elongation either by providing a mechanical barrier or by promoting peristaltic fluid movement inside the tubes (Jesudason et al., 2006). However, in kidneys, smooth muscles develop only next to the ureter and are not required for tube elongation (Bush et al., 2006). Similarly, in the mouse, biased outgrowth is already noted before airway smooth muscles (ASM) are first observed at E11.5 (Hines et al., 2013; Tang et al., 2011); peristaltic contractions are observed only from E12.5 (Schittny et al., 2000). Moreover, the inactivation of *Myocardin* prevents ASM differentiation, but does not prevent lung branching morphogenesis, and results in thinner, rather than wider branches (Young et al., 2020). Therefore, smooth muscles can be ruled out as a necessary driver of biased lung tube elongation. Importantly, branch shape is normal when *Mylk*, which encodes myosin light chain kinase (MLCK), an essential factor for ASM contraction, is inactivated during lung development (Young et al., 2020). This suggests that ASM cells may affect tube diameter via non-mechanical cues.

In this paper, we sought to systematically analyse the minimal requirements for biased epithelial tube elongation. To this end, we cultured mouse embryonic lungs and kidneys under different conditions and quantified the length and width of the branches for up to 60h. We show that the mesenchyme is not necessary as biased elongating outgrowth is still observed when epithelial buds are cultured on their own, in the absence of mesenchyme, embedded in an extracellular matrix (ECM) gel with uniformly dispersed growth factors. Furthermore, we show that while ERK signalling concentrates at the tip of branching isolated epithelial tubes, there is no evidence for the formation of actin-rich protrusions at the epithelial tips that could guide the biased elongating outgrowth. In early lung and kidney development, epithelial tubes only have a narrow luminal space, and tubular cross-sections are often elliptical rather than round. Despite the nonuniform curvature of such closed tubes, tension, as monitored with actin staining, remains uniform in the epithelium. We observe fluid flow inside the narrow embryonic tubes and show that the predicted shear stress level is within the range that cells can, in principle, sense. A cell-based model confirms that a tangential apical force, as provided by shear stress, can result in the reported bias in cell shape and elongating outgrowth.

## RESULTS

### Biased epithelial lung tube elongation

Given reports that the trachea switches from anisotropic to isotropic expansion around E12.5 (Kishimoto et al., 2018), we sought to measure the length and circumference of the bronchus of the left lobe (LL) between E10.5 and E14.5. For this, we used the Shh^GC/+^; ROSA^mT/mG^ transgenic mouse line, which expresses green fluorescent protein (GFP) in the cell membrane of the lung epithelium (Fig. 1A, C, D). Here, we averaged the circumference over the entire 3D bronchus, except for the parts where side branches form (Fig. S1). We confirm the previously reported 2-fold stronger longitudinal than circumferential expansion between E10.5 and E11.5 (Tang et al., 2011), and find that, much as in the trachea, there is a switch to isotropic growth at later stages, though a day later (E13.5) than in the trachea. The substantial widening of the bronchus thus occurs after the emergence of cartilage and smooth muscles (Hines et al., 2013; Schittny et al., 2000). Between E11.5 and E13.5, the bronchus still lengthens more than it widens, even though the overall rate of growth declines (Fig. 1E).

Each 3D length measurement in Fig. 1D,E comes from a different embryo, and we notice a certain level of variability between the specimen. Part of the differences can be accounted to differences in developmental progress, which is observed even in embryos from the same litter. To establish a reliable time line of the growth process, we cultured E11.5 embryonic lungs for 48h on a filter and measured the lengths and average diameter of the branches (Fig. 1F, Fig. S1). Given the development on a filter, there are differences in the branch angles, and much as for the 3D specimens, there is considerable variability between lungs. Nonetheless, in all specimens, we observe a similar biased expansion of the left bronchus (Fig. 1D,E, grey) as in the serially isolated embryonic lungs (Fig. 1D,E, green). The cultured lungs elongate slightly less than in the embryo, and there is less of a reduction in the branch width, though this difference may reflect differences in the analysis. The width in the 2D cultures was averaged along the entire branch (Fig. 1F), while the averaged circumference of the 3D specimen excluded the parts where branches emerge (Fig. S1). Overall, the cultured lungs recapitulate the growth process in the embryo very well, and we will therefore use these to analyse the mechanisms that drive elongating outgrowth.

### Mesenchyme is not required for biased epithelial tube elongation during lung and kidney development

While smooth muscles have recently been shown to be dispensable for lung branching morphogenesis (Young et al., 2020), and the lung and the kidney epithelium can grow and branch in the absence of the mesenchyme if an appropriate ECM gel and growth factors are provided (Nogawa and Ito, 1995; Qiao et al., 1999; Varner et al., 2015), the mesenchyme is well known to affect branch shapes (Blanc et al., 2012; Lin et al., 2003; Sakakura et al., 1976). To analyse the impact of the mesenchyme on biased epithelial outgrowth, we analysed the three lateral domain branches in the left bronchus (LL1-LL3), and two branches in the ureteric bud (UB) in cultured lungs and kidneys (Fig. 2A, C, Video S1, S2) as well as in explants, where the epithelium was enzymatically separated from the mesenchyme (Fig. 2E, G, Video S3, S4), embedded in Matrigel matrix, and suitable growth factors were added uniformly (Materials and Methods). We find that these five branches all decrease in their average diameter as they elongate (Fig. 2A-D). As a result, the elongation bias is even more pronounced than for the left lung bronchus. Both lung and ureteric buds still show biased elongating outgrowth when cultured in the absence of mesenchyme (Fig. 2F-H). This excludes a possible wall-like restrictive force, a pulling force, or other polarity cues from the mesenchyme as a necessary driver of epithelial tube elongation. It also confirms that smooth muscles are not necessary for biased elongating outgrowth. We note, however, that in the case of the ureteric bud, the branches elongate less and remain wider in the absence of mesenchyme. This shows that the mesenchyme impacts the elongation process, even though it is not necessary for biased elongation.

**Figure 2.**
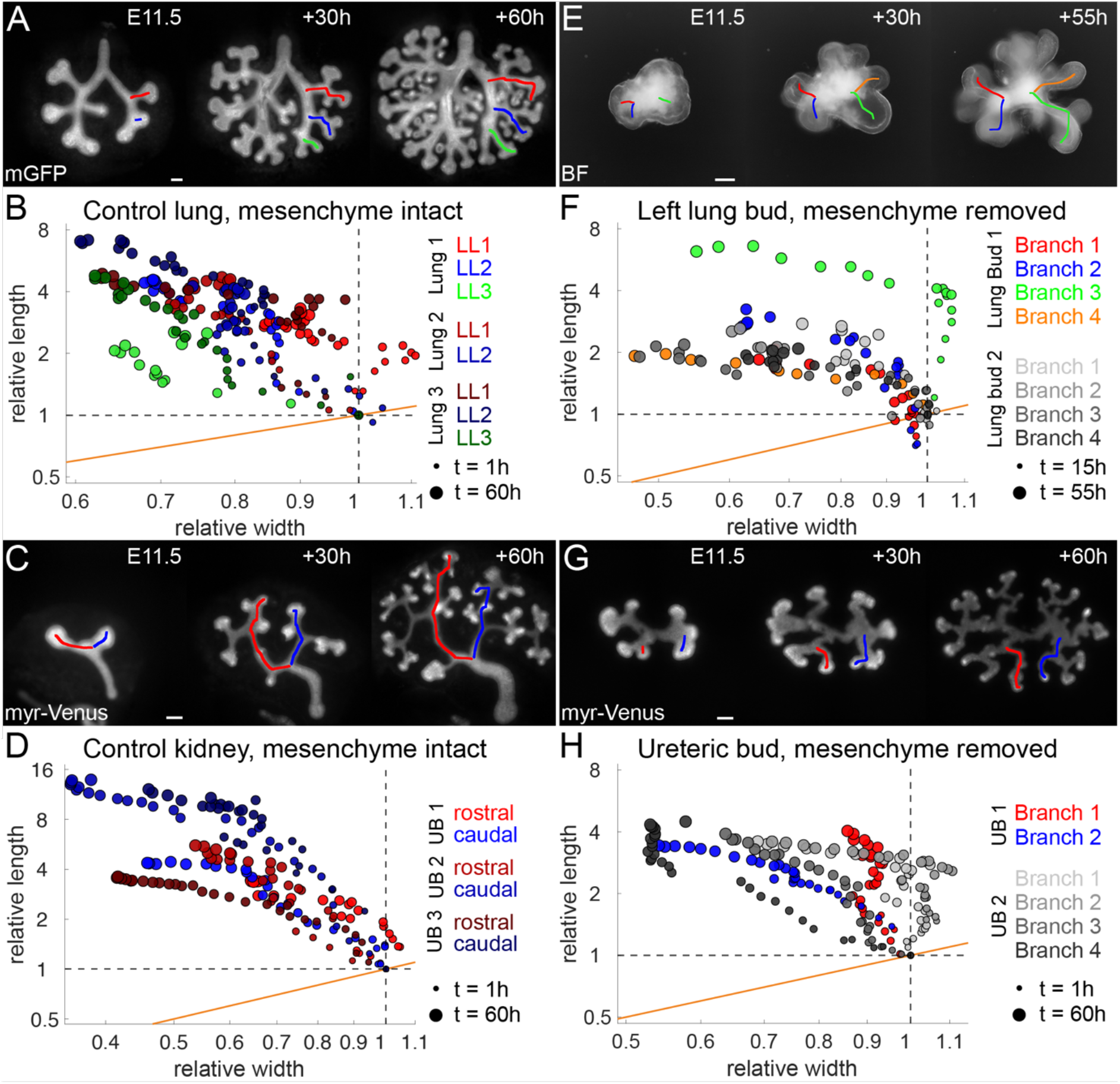
Mesenchyme is not required for biased epithelial tube elongation during lung and kidney development. **(A, C, E, G)** Epifluorescence/brightfield (inverted) microscopy images of the lung and kidney epithelium expressing mGFP or myr-Venus, respectively, and cultured for up to 60h. Samples in (E, G) are embedded in Matrigel (see materials and methods). Coloured lines mark the branches analysed in (B, D, F, H), respectively. Scale bars 100 μm. **(B, D, F, H)** Relative width and length measurements of lung and kidney epithelial branches. Data points above and away from the orange line show biased tube elongation, meaning branches elongate more than they widen. Dot size increases with culture time. **(A)** A representative control lung with intact mesenchyme. **(B)** LL1, LL2, and LL3 lung branches show biased epithelial tube elongation (sample n= branch n=8). **(C)** A representative control kidney with intact mesenchyme. **(D)** The rostral and caudal branches of the ureteric bud’s (UB) T-shape stage show a bias in tube elongation in control kidneys with intact mesenchyme (sample n=3; branch n=6). **(E)** A representative left lung bud epithelium without mesenchyme. **(F)** Without mesenchyme, lung branches continue to show biased elongation (sample n=2; branch n=8). The plotted time frame ranges from +15h until +55h, during which branches elongate (See Video S3). **(G)** A representative UB epithelium without mesenchyme. **(H)** Without mesenchyme, UB branches continue to show biased tube elongation. Compared to the control in (D), branches show a lesser relative increase in length and reduced branch thinning (sample n=3; branch n=6).

### Biased outgrowth is not driven by FGF signalling gradients

Signalling differs at the tips, i.e. phosphorylated ERK, which signals downstream of FGF10/GDNF (Fig. 3A) and which is necessary for lung branching morphogenesis (Boucherat et al., 2014), localizes at the tips in embryonic lungs (Fig. 3B), kidneys (Fig. 3C), as well as in mesenchyme-free cultures (Fig. 3D). Accordingly, a pulling force from the tip could, in principle, drive the uniform biased outgrowth along the entire tube (Fig. 3E). However, no actin-rich protrusions, such as filopodia or lamellipodia, are observed at the basal epithelial tips. The staining for actin is instead strongest along the apical luminal surfaces (Fig. 3F, G, arrows). Even though FGF10 can act as chemoattractant for explant lungs (Park et al., 1998), and the FGF receptor (FGFR) inhibitor SU5402 reduces the overall growth rate (Goodwin et al., 2019), SU5402 does not reduce the elongation bias (Fig. 3H, Fig. S2, Video S5). This is consistent with previous observations that lung epithelial tubes continue to elongate in *Fgf10* and *Fgfr2* conditional mutants (Abler et al., 2009), but is inconsistent with a role for FGF signalling as a driver of biased elongating outgrowth.

**Figure 3.**
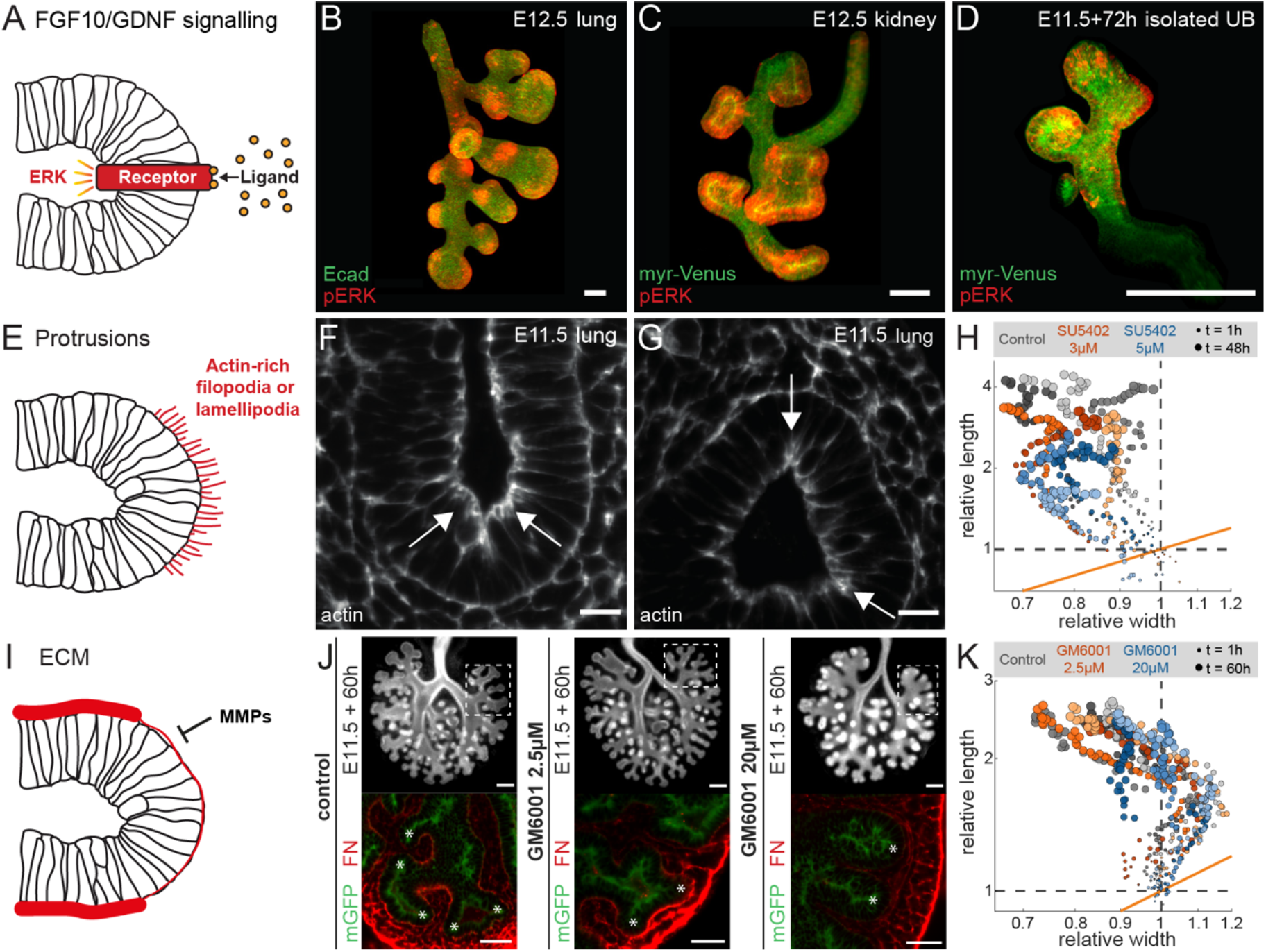
Biased outgrowth is not the result of FGF signalling or ECM turn-over at the tip. **(A)** Concentrated growth factor signalling might lead to biased outgrowth. **(B-D)** pERK antibody staining shows localized spots at the tips of **(B)** an E12.5 embryonic lung, **(C)** an E12.5 kidney, and **(D)** a cultured isolated ureteric bud (UB). Scale bars 100 μm. **(E)** A pulling force due to cytoskeletal protrusions could drive biased outgrowth. **(F,G)** Actin staining of E11.5 embryonic lungs shows enrichment at the apical, but not at the basal tissue boundary. Arrows point to regions of high actin intensity. Scale bars 20 μm. **(H)** Relative width and length measurements for the first secondary branch of the left lobe (LL1) of E11.5 control and FGFR inhibitor-treated lungs over 48h. Different colour shades distinguish the data for different lung samples (n=3 for each condition). **(I)** ECM degradation by MMPs at the tip could drive biased outgrowth. **(J)** Epifluorescence microscopy images of E11.5 control and MMP inhibitor-treated lung epithelium, expressing mGFP, after culture on a filter for 60h (top row). Dashed boxes mark the region of LL1 branches. Scale bars 200 μm. Fibronectin (FN) antibody staining of LL1 branches (bottom row) shows low intensity at the tips, but high intensity in cleft regions of control lungs. Lungs treated with 2.5 μM MMP inhibitor have high fibronectin intensity in both tip and cleft regions. In lungs treated with 20 μM MMP inhibitor, fibronectin is deposited between mesenchymal cells rather than in the basement membrane. Asteriks mark branch tips. Scale bars 50 μm. **(K)** Relative width and length measurements for the first secondary branch of the left lobe (LL1) of E11.5 control and MMP inhibitor-treated lungs over 60h. Different colour shades distinguish the data for different lung samples (n=3 for each condition).

### Biased outgrowth is not the result of ECM thinning at the tip

Matrix metalloproteinase (MMP) support lung bud outgrowth by degrading ECM preferentially at the lung tips such that the basement membrane, composed of nidogen, collagen, fibronectin, and laminin, is thinner at the lung tips (Fig. 3I,J) (Gill et al., 2003; Gill et al., 2006; Mollard and Dziadek, 1998; Moore et al., 2005; Rutledge et al., 2019). The MMP inhibitor GM6001 interferes with ECM digestion and enhances lung growth and branching at low concentrations, but impairs outgrowth at high concentrations (Gill et al., 2006) (Fig. 3J). Despite its impact on lung growth and branching, neither low (2.5 μM) nor high (20 μM) GM6001 concentrations affect the bias of tube outgrowth in E11.5 explant cultures (Fig. 3K, Fig. S3, Video S6).

### Mechanically forced tube collapse does not result in directional cues for uniform biased outgrowth

Cross-sections of a mouse embryonic lung reveal elliptical tissue shapes with closely opposing apical sides and narrow luminal spaces (Fig. 4A, yellow outlines). Wider lumina are only observed close to branch points and in the tips (Fig. 4A, blue outlines). Collapsed tubular morphologies are not the result of either tissue processing or clearing, as they are also observed in developing lungs cultured in a light-sheet microscope for over 34h (Fig. 4B, Fig. S4, Video S7). Similar observations are made in embryonic kidneys (Fig. 4C).

**Figure 4.**
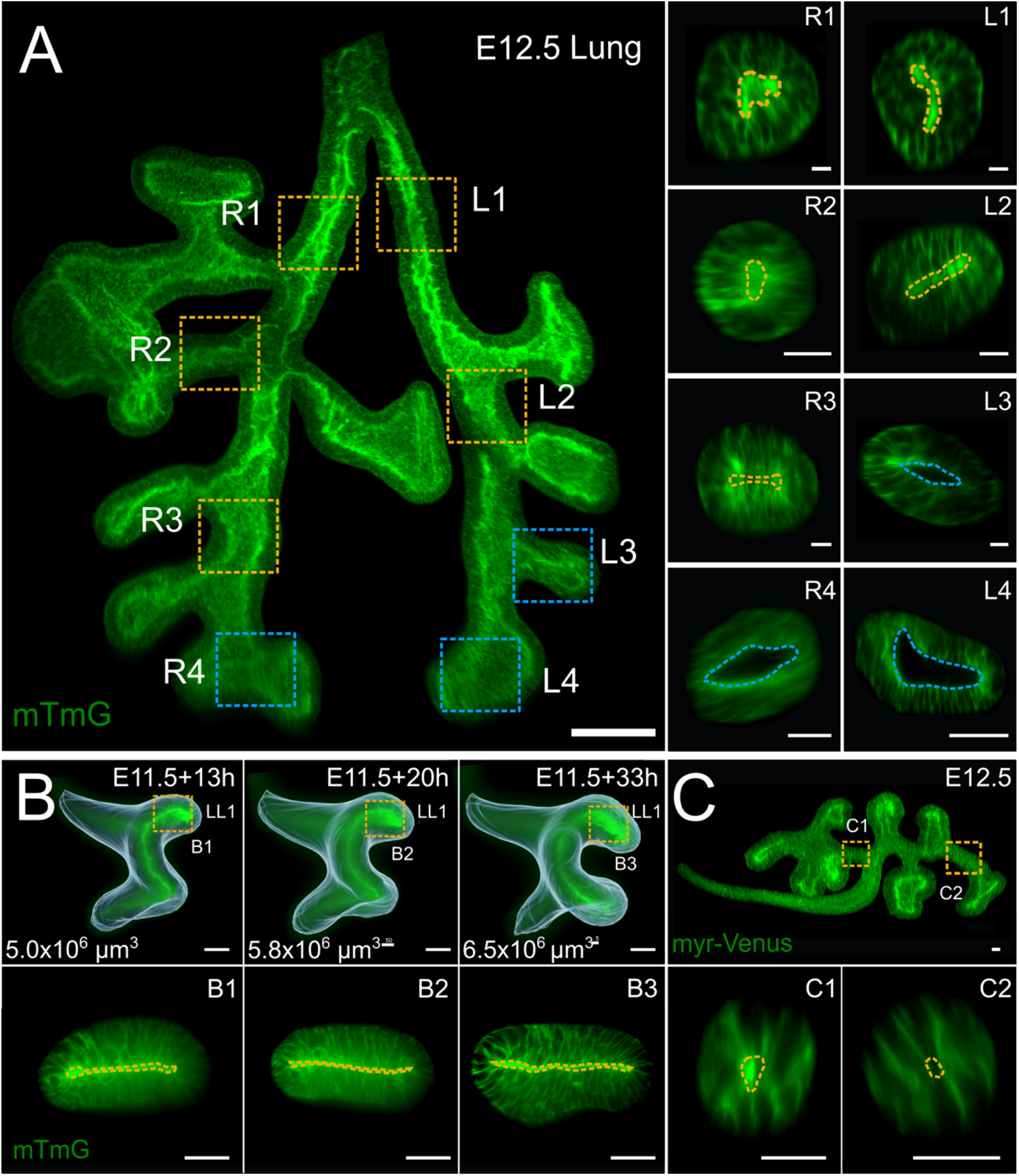
Collapsed epithelial tubes in embryonic mouse lungs and kidneys. **(A)** Light-sheet fluorescence microscopy image of CUBIC-cleared embryonic lung. Dotted boxes mark different regions in the branching tree where luminal morphology was analysed. Cross-sections illustrate elliptical tissue shapes with narrow luminal spaces in tubular sections (yellow outlines), as well as wider luminal spaces in tips and branch points (blue outlines). Scale bars 50 μm. **(B)** High-resolution light-sheet microscopy time-lapse imaging of embryonic lung development. Isosurface overlays highlight shape changes; overall domain volumes are given on the lower left. Cross-sections in boxed regions (lower panels) corroborate dynamic collapsed morphologies in elongating epithelial tubes (yellow outlines). The specimen was imaged for over 40h every 20min. Scale bars 50 μm. **(C)** Embryonic kidney epithelia also display narrowed tubular architectures (yellow outlines). Scale bars 25 μm.

To explore the possible mechanical effects that can lead to the observed collapse of epithelial tubes, and whether this could provide cues for uniform biased tube outgrowth, we conducted continuum-mechanical finite element simulations (Fig. 5A). Since tube collapse is the same along the tube length up to minor boundary effects for a cylindrical geometry, we can exploit the symmetry by considering only a two-dimensional cross-section perpendicular to the tube axis. In our numerical model, the tubular epithelial tissue was represented by an isotropic, linearly viscoelastic continuum, neglecting the cellular structure of the tissue. The epithelial material properties were therefore characterized by a Young modulus *E* and a Poisson ratio *ν*. As an initial condition, we chose a tubular shape with uniform radius *R* (measured from the cylinder axis to the middle of the tissue), and the relative tissue thickness was set to *t/R=0.5*. The epithelium was set to be intrinsically uncurved, such that a stress-free configuration would be a flat tissue. We used a custom-built finite element simulation framework developed for large deformations of thin (visco-)elastic structures (Vetter et al., 2013). For simulation details, see Materials and Methods.

**Figure 5.**
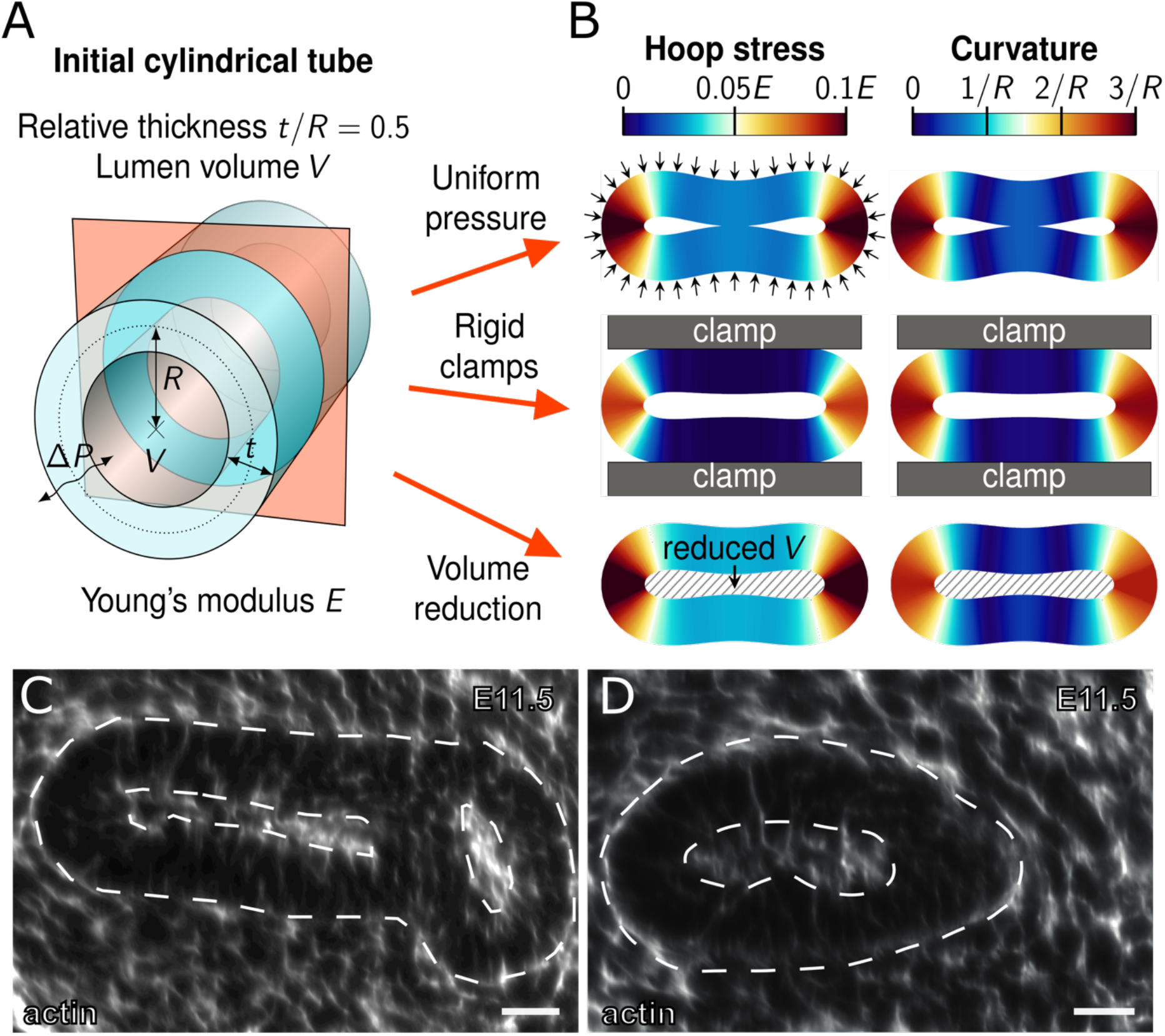
Mechanically forced tube collapse does not result in directional cues for uniform biased outgrowth. **(A,B)** Mechanical finite element simulations of collapsing epithelial tubes. **(A)** Schematic of the initial tubular configuration. A two-dimensional cross-section (red) was simulated using linear viscoelasticity. **(B)** Resulting collapsed shapes for three different scenarios: a uniform pressure difference *ΔP* (top row, arrows), rigid external clamps (middle row, grey bars), and a reduced lumen volume *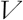* (bottom row, hatched area). Colours indicate the measured hoop stress (left column) in units of Young’s modulus *E*, and the midline curvature (right column) in units of the inverse initial tube radius, *1/R*. Hoop stress is localised almost exclusively in the highly curved extremal regions. **(C,D)** Actin staining is higher at the apical side, but otherwise uniform in cross-sections of closed epithelial tubes from an E11.5 mouse lung. Scale bars 30 μm.

We considered three different collapse scenarios. In the first, a uniform net pressure difference *ΔP* was applied, corresponding to either a pressure drop in the lumen or an increased pressure exerted onto the epithelium by the external environment. The pressure was increased until the critical point of collapse was surpassed. In the second scenario, the epithelial tube was pinched by two rigid parallel clamps slowly approaching one another, mimicking external spatial constraints imposed by a stiff surrounding medium. In the third scenario, the enclosed lumen volume 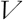 was controlled with a Lagrange multiplier and drained over time until the tube was sufficiently collapsed. Figure 5B shows the equilibrated simulation results for each of the three scenarios. In all cases, both the hoop stress and curvature profiles along the tissue midline are highly nonuniform. Hoop stress is localized almost exclusively in the two extremal points with large curvature. Extruded along the tube axis, these point-like localisations form tangential lines in the longitudinal direction, which could, in principle, provide the required directionality. However, any driving force for straight tube outgrowth would be expected to act relatively uniformly along the circumference. Without such uniformity, biased outgrowth would lead to curving of growing tubular segments or other distortions in the growth pattern, also on the cellular level, which are not generally observed. We conclude that, given their nonuniform distribution in the tube cross-section, the stress and curvature patterns that arise from the deformation cannot serve as cues for uniform biased outgrowth.

Stresses can be relaxed rapidly in tissues. We sought experimental evidence for residual stress or increased tension in the most curved parts of the epithelium. Actin has been reported to be able to function as a tension sensor (Hayakawa et al., 2011) and to polymerise under external mechanical stimuli (Hirata et al., 2008; Shao et al., 2015). From antibody staining for actin, we found a relatively uniform actin density in the closed lung tubes (Fig. 5C,D). While this does not conclusively rule out the possibility of residual stress patterns, we found no evidence for it. It remains possible that a wall-like constraint, in combination with rapid stress relaxation, enforces the elliptic shape and elongating outgrowth. However, it is unclear how such an outer wall-like constricting force would arise even in the absence of mesenchyme. Moreover, we have shown before that a constricting force that results in the observed biased epithelial outgrowth in a cell-based model is insufficient to generate the observed bias in cell shape and cell division (Stopka et al., 2019). Consequently, the mechanical constraints explored here are unlikely to drive the biased elongating outgrowth of embryonic lung tubes.

### Shear stress in the developing lung

In later stages of lung and kidney development, there is evidence for fluid flow inside the epithelial tubes (Blewett et al., 1996; Harding et al., 1984; Nelson et al., 2017; Unbekandt et al., 2008; Vasilyev and Drummond, 2014; Vasilyev et al., 2009). Fluid flow results in shear forces that act tangentially on the apical side of the epithelial cells, and could thus, in principle, drive elongating outgrowth (Fig. 6A). The flow rate has previously been estimated as 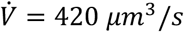 in developing lungs (George et al., 2015), but this value was inferred based on measurements at fetal stage in other species than mouse. We, therefore, developed a protocol to measure the flow velocity in cultured E11.5 mouse lung rudiments. To this end, we injected fluorescent beads into freshly dissected E11.5 lung rudiments, transferred them into a low-volume culture system, adapted from (Sebinger et al., 2010), and waited for at least 30 minutes to allow the injected lungs to recover, and imaged the developing lungs for up to 2 hours under a spinning disk confocal microscope (Fig. 6B, Fig. S5, Video S8). The fluid velocity is highest in the center of the tube and zero at the walls (Fig. 6A). As we cannot image the lung in 3D, we cannot determine the distance of the beads to the wall. Accordingly, we do not know how far the fastest-moving beads are away from the center. By focussing on the fastest-moving beads in five independent injection experiments, we thus obtain a lower bound on the maximal flow velocity of *u =* 0.73 ± 0.26 *μm/s* (n=5). We emphasize that this provides a conservative estimate of the flow velocity and thus the shear stress estimate. The true fluid velocity and shear stress may still be larger.

**Figure 6.**
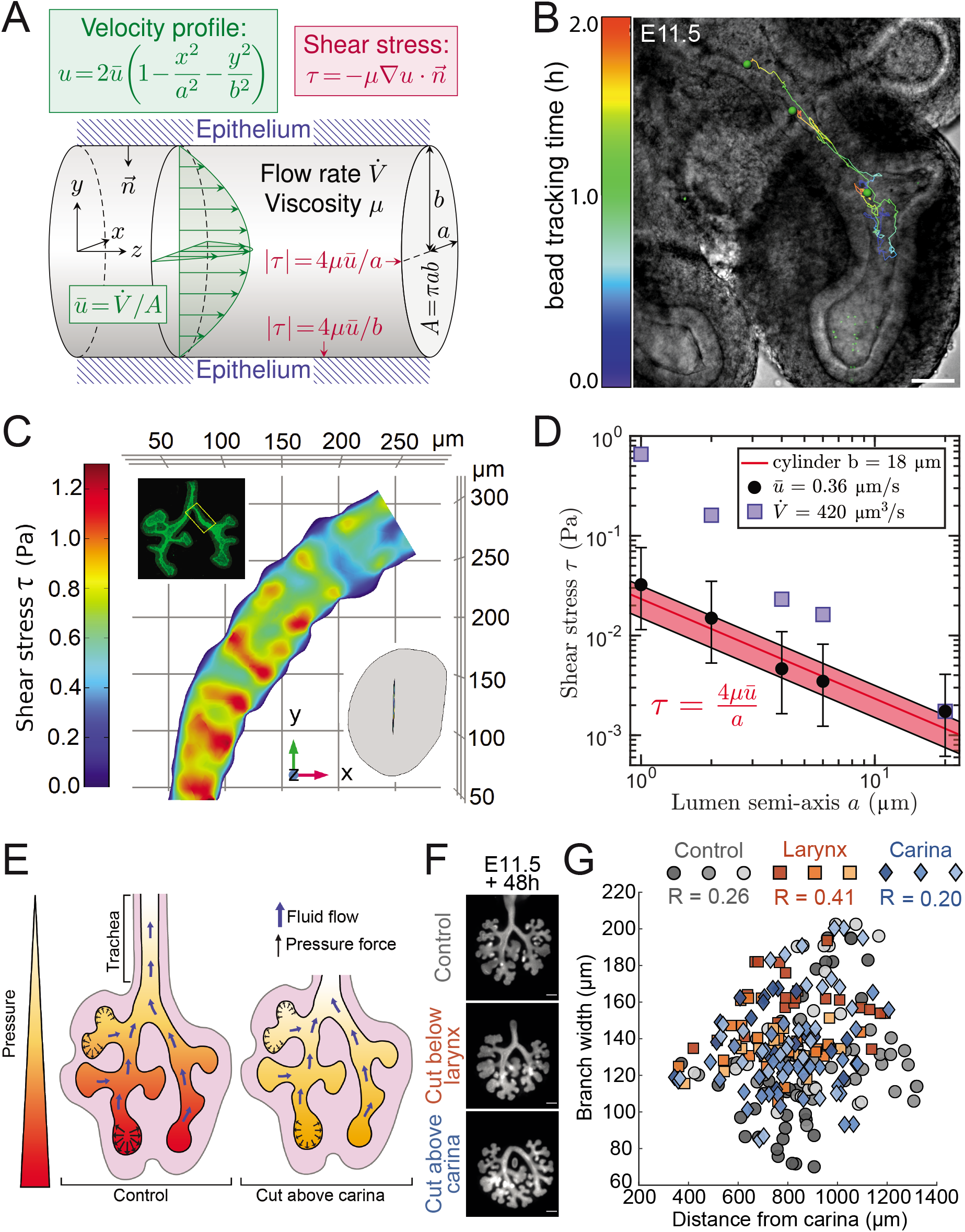
Shear stress in the developing lung. **(A)** Schematic of shear stress for Hagen-Poiseuille flow in a tubular lumen with elliptical cross section. Wall shear stress | *τ*| is maximal in weakly curved regions (minor ellipse axis) and minimal in strongly curved regions (major ellipse axis) of the luminal surface. **(B)** Beads were injected into the lumen of the distal-most tip of an E11.5 lung and imaged over 2 h. Composite of a brightfield image of the right lung lobe and the confocal channel (488 nm excitation) of the beads, overlayed with the bead tracks (generated using Imaris Track, Autoregressive motion algorithm). The tracks that show directional movement towards the trachea are displayed (representative image, n = 5). Scale bar 70 μm. **(C)** Estimated wall shear stress levels at the apical surface of a 400 μm epithelial tube segment that was extracted between the carina and the first branch of the left lung lobe (LL1) (inset). The lumen geometry after numerical rescaling had an average cross sectional opening of 2 μm. **(D)** The lumen geometry presented in Fig 6B was rescaled with different scaling factors. For each scaling factor employed, the semi-axis a of the lumen was measured. We then calculated the average wall shear stress on the apical surface τ induced by a flow with an average velocity of 0.36 μm/s (black dots) or a flow rate of 420 μm^3^/s (blue squares). The error bars correspond to the standard deviation of the average wall shear stress. The average level of wall shear stress for a given flow velocity can be well approximated from a Hagen-Poiseuille flow profile in an elliptical tube of equivalent size (red line). The red shaded region was determined with the same approximation for a flow with an average velocity ± the measured standard deviation of 0.13 μm/s. **(E)** Fluid secreted by the cells at the bud tips flows toward the trachea opening. For such a flow to exist, the fluid pressure must be higher at the tips than at the trachea opening. In the case of a FSI, the luminal fluid pressure would widen the branches at the tips more than the branches near the trachea. The existence of such FSI can be tested by reducing the pressure difference between the tips and the outlet. This was achieved by culturing the lungs without their trachea. **(F)** Epifluorescence microscopy images of E11.5 lung epithelium, expressing mGFP, after culture on a filter for 48h. The distance of the branches from the tracheal opening was altered by cutting the trachea either below the larynx or above the carina prior to the culture. Control lungs maintained an intact trachea. Scale bars 200 μm. **(G)** Branch width and distance measurements for all branches of the left lobe of the lung cultures in (E) at the culture endpoint. Different symbol types mark the different culture conditions regarding trachea length. Different colour shades distinguish the data for different lung samples (n = 3 for each condition). Branch width does not depend on the distance from the tracheal opening, reflected by low Pearson correlation coefficients R.

The shear stress level depends on the flow velocity, the viscosity of the fluid, and the shape and size of the tube lumen (Fig. 6A). The viscosity of the fluid in the lung lumen has been measured to be 10 times higher than that of water, i.e. *μ =* 0.016 *Pa · s* (Bokka et al., 2015). To evaluate the shear stress levels on the apical side of embryonic lungs, we simulated Stokes flow through the lung tube of an E11.5 embryo (Fig. 6C). The lung tube geometry was obtained by segmenting a 400 μm section between the carina and the first branch of the left lung lobe of an E11.5 lung (Fig. 6C, top inset). The average luminal width is about 2 μm (Fig. 6C, bottom inset). We note that given the strong fluorescence on the apical side, the segmentation was possible only in subapical layer. Accordingly, the extracted geometry had a luminal minor axis of 20 μm, which we computationally reduced to the measured luminal minor axis of 2 μm. With these values, we find that the shear stress is highest on the flat part of the embryonic lung tube, and mostly in the range 0.01-0.1 Pa (Fig. 6D, black circles). As we computationally alter the luminal width, we find that the average level of shear stress on the apical surface can be well approximated from a Hagen-Poiseuille flow profile in an elliptical tube of equivalent size (Fig. 6A, D, red line). With the original extracted shape (20 μm along its minor axis), the predicted shear stress level is therefore about 10^-3^ Pa. We note that impressively similar shear stress levels are predicted with the previously estimated volume flow rate, 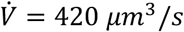 (Fig. 6D, blue squares).

Shear stress does not directly deform cells, but rather cells sense shear stress via their primary cilium and actively respond with cell shape changes (Galbraith et al., 1998; Weinbaum et al., 2011). Epithelial kidney cells are particularly sensitive, with renal collecting duct chief cells responding to apical shear stress as low as 6.8·10^-4^ Pa (Resnick and Hopfer, 2007), and cultured epithelial kidney epithelial cells responding to 0.075 Pa, but not to 1.5 Pa (Nauli et al., 2003). On the other end of the spectrum, endothelial cells are particularly insensitive and typically respond to 0.5-5 Pa (Weinbaum et al., 2011). Cornelial epithelial cells respond to 0.4-0.8 Pa (Molladavoodi et al., 2017), while alveolar epithelial cells respond to 0.7-1.5 Pa (Flitney et al., 2009). While primary cilia have been reported in lung epithelial cells (Jain et al., 2010), the sensitivity of epithelial cells in mouse embryonic lungs and kidneys has not yet been reported, but is likely in a similar range. Accordingly, cells should be able to sense apical shear stress with the measured luminal width. Consistent with a role of shear stress in driving elongating outgrowth, in embryonic lungs that express the constitutively active form KRas^G12D^ in the epithelium (Shh^cre/+;^KRas^LSL-G12D/+^), which disrupts the primary cilium (Tape et al., 2016), the bias in cell division and epithelial outgrowth is lost (Tang et al., 2011).

The pressure gradient in the lumen could also, in principle, impact on tube width directly through a fluid-structure interaction (FSI). To obtain a flow from the bud tips to the opening of the trachea and the fluid outflow from the trachea that is observed in cultured explants (Video S9), the fluid pressure must be highest at the tip and smallest at the tracheal opening (Fig. 6E). In case of a FSI, the shape of the branches would then depend on the local fluid pressure, and buds should be wider than stalks. While stalks are indeed thinner than buds, there is no direct dependency of branch width on the distance from the tracheal opening (Fig. 6F, Fig. S6, Video S10). A simple way to modulate the pressure at the tips is by altering the distance between the tips and the outlet by culturing lungs either with or without their trachea (Fig. 6F). Removal of the trachea shortens the distance to the outlet, and thus, in case of a constant pressure gradient and flow rate, reduces the pressure difference between the tips and the outlet. We find, however, that a removal of the trachea neither impacts branching morphogenesis nor tip shapes (Fig. 6F,G), which rules out a significant mechanical impact of the fluid pressure on the surrounding epithelium.

### Forces in the longitudinal direction can result in the observed bias in cell shape and outgrowth

Finally, we used a cell-based tissue model to investigate whether a biased force in the longitudinal direction, as arises from shear stress, could recapitulate the observed bias in the apical cell shapes, cell division, and elongating outgrowth (Fig. 7A). In the cell-based tissue model, it does not matter whether the force that deforms the apical cell surface arises directly from the shear stress acting directly on the cell or indirectly via the intracellular response of the cell to shear stress. The intracellular response to shear stress only means that the absolute force levels that deform the cell do not need to correspond to the shear stress level that is sensed by the cells. Accordingly, we will not attempt to relate the force levels in the simulations to the estimated shear stress levels above, and rather use arbitrary units (a.u.).

**Figure 7.**
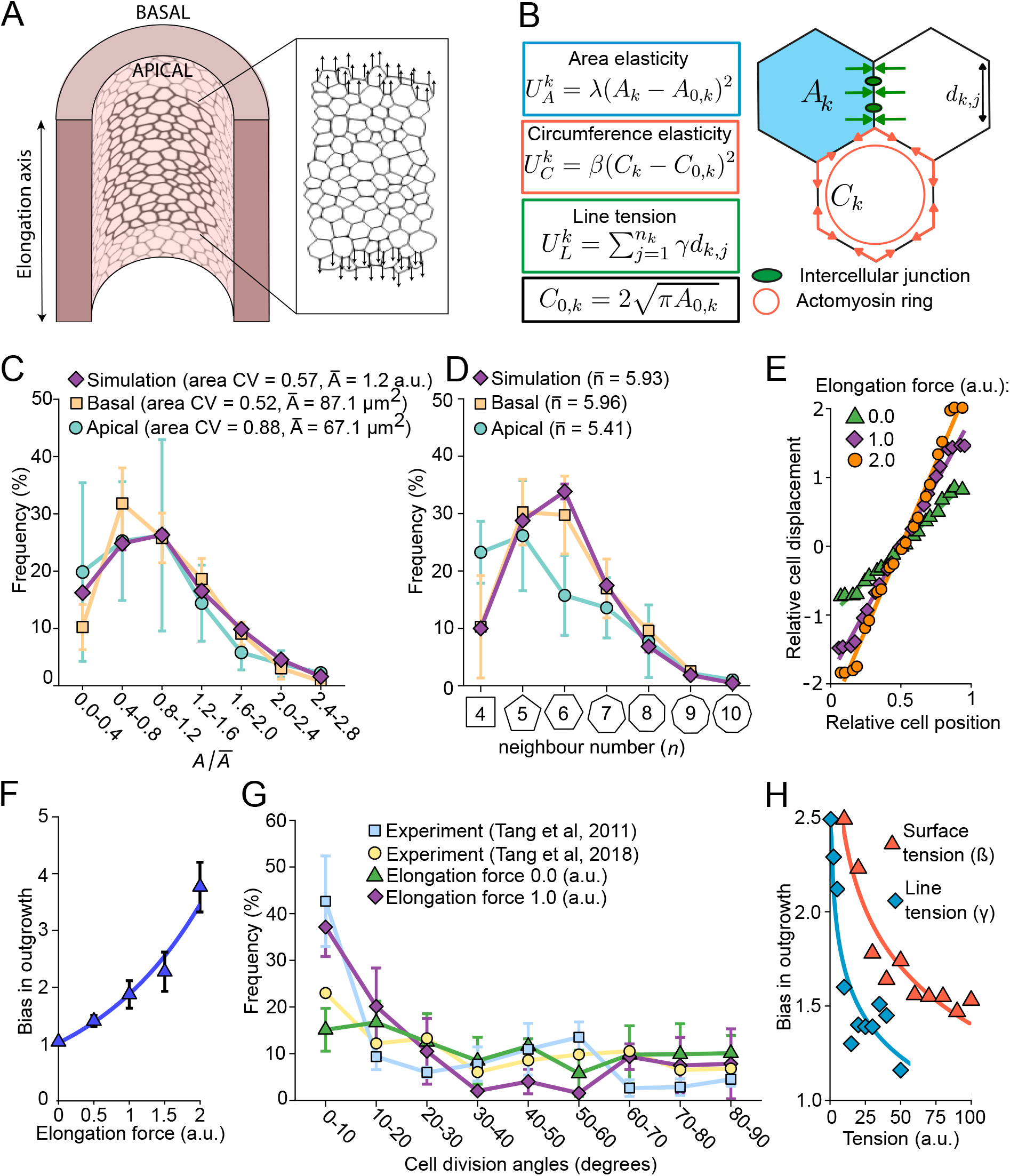
Longitudinally biased forces can result in the observed bias in cell shape and outgrowth. **(A)** The apical surface of the lung epithelium was simulated with a 2D vertex model. Longitudinally biased external forces were applied to the vertices of the cells at the top-most and bottom-most layers of the tissue. **(B)** Schematic of the cell-based model. The potential energy of the system is comprised of three contributions. The area elasticity energy *U_A_* penalizes any deviation of the cell area Ak from its target area *A*_0, *k*_ The constant *λ* defines how resistant the cells are to deformations. Similarly, the circumference elasticity energy *U_c_* aims to emulate the contractility of the actomyosin ring by penalizing any deviation of the cell circumference *C_k_* from its target circumference *C*_0,*k*_. The line tension *U_L_* energy gives rise to a force associated with cell-cell adhesion. Low values of *γ* characterize stable and favourable contacts between cells. **(C,D)** The cell area (C) and cell shape (D) distributions of E11.5 lung epithelial cells were measured at their apical (green) and basal (yellow) sides. All parameters in the vertex model (purple) were set so that simulated tissues reproduce the measured distributions (n = 3 for experimental tissues and n = 5 for simulated tissues, error bars show standard deviations). **(E)** Relative displacement of the cells as a function of their initial positions in the simulated tissues. Regardless of the magnitude of the elongation force applied, the cells are displaced uniformly along the tissue axis during the simulations. **(F)** Bias in outgrowth of the simulated tissues as a function of the elongation forces applied. A force of 1.0 a.u. yields the measured (Tang et al., 2011; Tang et al., 2018) 2-fold elongation bias of lung tubes (n = 5, error bars show standard deviations). **(G)** Distribution of the cell division angles in the simulated tissues in the presence or absence of an elongation force. The elongation force 1.0 a.u. yields a bias in cell division orientation equivalent to the bias reported by (Tang et al., 2011); n = 5 for simulated tissues, error bars show standard deviations. **(H)** Bias in outgrowth of tissues subjected to an elongation force of 1.5 a.u. as a function of their surface tensions or line tensions. High cortical tensions result in reduced biased outgrowth.

We implemented a 2D vertex model (Fig. 7B) using the open-source software Chaste (Fletcher et al., 2013; Mirams et al., 2013). The model has eight biophysical parameters (Supplementary Table 2). With the standard parameters (Fletcher et al., 2013), we can reproduce the characteristics of polygonal epithelial lattices (Fig. S7) (Kokic et al., 2019; Vetter et al., 2019). Epithelia differ in their area and cell shape distributions. We, therefore, used light-sheet microscopy to determine the apical and basal cell area and cell shape distributions in an E11.5 lung tube (Fig. 7C, D). To reproduce the wide area distributions, we had to introduce a Gaussian distribution for the cell cycle times and the cell division thresholds (Supplementary Table 2) (Kokic et al., 2019). For the sake of simplicity and given the absence of contrary data, we model the effect of shear stress in our cell-based tissue model by applying a force at one tissue border in the direction of the shear stress (Fig. 7A). This results in a uniform force field with a uniform relative displacement of cells along the tissue axis (Fig. 7E, Video S11), as would be expected in the case of shear stress. When we apply uniform growth, we find an almost linear increase of the bias in outgrowth with such an external force (Fig. 7E,F).

Between E10.5 and E11.5, the lung tubes elongate twice more than they widen (Fig. 1E) (Tang et al., 2011). We obtain this 2-fold bias in outgrowth with an elongation force of 1 a.u. (Fig. 7F). It has previously been noted that this bias in outgrowth is accompanied by a bias in cell shape and cell division (Tang et al., 2011; Tang et al., 2018). Cell shape and the cell division axis are linked in that cells in the lung epithelium divide perpendicular to their longest axis when their aspect ratios are greater than 1.53 (Tang et al., 2018). With a force of 1 a.u. and cell division perpendicular to the longest axis, the simulations (Fig. 7G, purple diamonds) recapitulate the measured bias in cell division at E10.5 (Fig. 7G, blue squares, (Tang et al., 2011)), as well as our measured cell area and shape variation (Fig. 7C,D). Note that a slightly lower bias in the cell division angles has been reported at E11.5 (Fig. 7G, yellow circles, (Tang et al., 2018)). We also recapitulate the observed impact of additional stretching forces (Tang et al., 2018) on elongating outgrowth (Fig. 7E,F). Finally, we note that the longitudinally biased force on elongating outgrowth depends on the mechanical properties of the cells in our simulations (Fig. 6H). In particular, the higher the cortical or line tension, the smaller the bias, and *vice versa*. The changes in the elongation bias in different mutants and pharmacological perturbations may thus also arise via modulation of cortical tension and cell-cell adhesion. Given that such parameter changes can also affect the cell area and shape variability (Fig. S7), we note that the simultaneous recapitulation of the elongation bias (Fig. 7F), and the cell area and shape distributions (Fig. 7C,D) is significant. We conclude that a force that is biased in the longitudinal direction allows us to recapitulate all measured features.

## DISCUSSION

The elongation of epithelial tubes is a key developmental process. We combined a quantitative analysis of lung and kidney branching morphogenesis with computational modelling to evaluate candidate mechanisms for the biased elongation of epithelial tubes. We show that biased elongation is an inherent property of these epithelial tubes, and that it does not require contact with the mesenchyme or an external chemotactic gradient. We note that the epithelial tubes are largely collapsed in early lung and kidney development, and we show that fluid flow exists already early in lung development. The measured flow velocity is sufficiently high that the resulting shear stress can be sensed by epithelial cells with their primary cilium (Weinbaum et al., 2011). We evaluate the impact of shear stress in a cell-based tissue model, and find that shear stress, unlike constricting forces (Stopka et al., 2019), can explain both the observed biased tube elongation and the observed bias in cell division. Shear stress may thus be a more general driver of biased tube elongation beyond its established role in angiogenesis (Davies, 2009; Galbraith et al., 1998; Galie et al., 2014).

Consistent with a role for shear stress in biased lung tube elongation, the bias in cell division and outgrowth is not observed in embryonic lung epithelia that express the constitutively active form KRas^G12D^ (Tang et al., 2011), a mutation that leads to the disruption of the primary cilium (Tape et al., 2016). The cilium is necessary to respond to shear stress because shear stress does not directly deform cells, but rather cells sense shear stress via their primary cilium and actively respond with cell shape changes (Galbraith et al., 1998; Weinbaum et al., 2011). Cells then divide perpendicular to their longest axis (Hertwig, 1884), and the bias in cell shape along the lung tube axis therefore translates into a bias in cell division (Tang et al., 2011; Tang et al., 2018).

Given that shear stress is actively sensed and translated by the cells into a change in cell shape (Galbraith et al., 1998), the force that deforms the cell does not need to correspond to the shear stress level that is sensed by the cells. With a force that lets us quantitatively recapitulate the measured bias in outgrowth, we also quantitatively reproduce the measured distribution in cell division angles as observed in the mouse lung bud (Tang et al., 2011; Tang et al., 2018). Importantly, in the embryonic mouse lung cells, cells only follow this longest-axis rule if the ratio of the longest to the shortest axis is larger than 1.5; otherwise, the cell division axis is set randomly (Tang et al., 2018). We emphasize that we can reproduce the measured elongation bias and the measured distribution of cell division angles only with the line tension and cortical tension parameters that allow us to also recapitulate the measured apical area and cell neighbour distributions. Smaller tension levels result in more elongation and *vice versa*. Accordingly, mutations that change the epithelial cell mechanics or the composition of the extracellular matrix (ECM) would be expected to affect the elongation bias in tube outgrowth. In addition to its effect on the primary cilium, KRas has previously also been linked to changes in cell shape and motility in airway epithelial cells by affecting cortical actin (Fotiadou et al., 2007; Okudela et al., 2009), and the KRas^G12D^ mutation has been found to upregulate multiple ECM components in the pancreatic stroma (Tape et al., 2016).

While biased tube elongation is observed in isolation, independent of the mesenchyme, we find that the mesenchyme affects the strength of this bias. In this way, the mesenchyme can modulate the particular tube shape and thereby the organ-specific branching architecture (Lin et al., 2003). It is not known how the mesenchyme impacts on lung tube elongation. However, several soluble factors that are secreted in the mesenchyme are known to signal in the epithelium and to affect the composition of the ECM (McCulley et al., 2015). Smooth muscles and the peristaltic fluid movement inside the lung tubes that they generate are not necessary for normal lung branching morphogenesis (Young et al., 2020). We can also exclude that the luminal pressure has a significant impact on tube shape. If the flow was to directly mechanically affect the epithelial cells, one would expect both a stretching of the cells in direction of the shear stress and a widening of the tube in response to the local luminal pressure. As flow is the result of a pressure gradient, the circumference of the tubes would then need to be wider with increasing distance from the larynx. This is not the case. Also, branching morphogenesis of cultured lungs is not affected by the presence or absence of the trachea. Accordingly, a fluid-structure interaction appears to have no significant impact on lung tube shape.

Shear stress only has the potential to drive biased elongating outgrowth because the epithelial tubes are so narrow in early lung and kidney development. In later stages, tubes are wide and open. The same level of apical shear stress would then require much higher flow rates, and tube growth indeed becomes isotropic. It remains unclear why the tubes collapse in early developmental stages. Mechanical effects that could, in principle, cause the collapse of the tubes would result in the highest mechanical stress levels in the curved parts. Accordingly, neither curvature nor hoop stress (Hamant et al., 2008) could explain the biased uniform outgrowth of the collapsed tubes. We note that staining for actin, a read-out for tension within tissues, is uniform in the closed tubes, suggesting that any stress that may have been generated during the collapse is quickly relaxed away. Going forward it will be important to identify the cause for tube collapse and understand its potential impact in biasing elongating outgrowth.

Beyond its role in driving elongating outgrowth in early lung development, fluid flow could also play a role in the reorganisation of the epithelial tree into a fractal-like architecture at later stages of development (Iber and Menshykau, 2013). Given that shear stress is actively sensed and translated by the cells into a change in cell shape (Galbraith et al., 1998), there is, however, not necessarily a linear correspondence between the extracellular shear force and the intracellular force that reshapes the cell. Other force response curves could result from the intracellular regulatory processes that respond to the shear stress. Microfluidic experiments could be used to study this aspect quantitatively. Such setups have previously been used to analyse the impact of the pressure differences between the larynx and the pleural space on the growth and branching behaviour with more precision that had been possible with surgical occlusion experiments (Blewett et al., 1996; Nelson et al., 2017; Unbekandt et al., 2008). The branch shapes have not been quantified in these studies, but appear shorter, pointing to a reduced elongation bias, which would be consistent with the expected reduced fluid outflow.

In summary, by combining quantitative measurements with simulations, we have shown that flow-induced shear stress can quantitatively explain the biased elongating outgrowth of epithelial tubes in early lung and kidney development. Going forward, it will be interesting to understand how the mesenchyme modulates the tube elongation bias to achieve organ-specific tube shapes, and whether fluid flow plays a role in generating the fractal-like tree architecture that emerges later in lung development.

## MATERIALS AND METHODS

### Ethical Statement

All experiments were performed in accordance with Swiss federal law and the ordinance provided by the Canton Basel-Stadt and approved by the veterinary office of the Canton Basel-Stadt, Switzerland (approval number 2777/26711).

### Mouse strains

Mice were housed at the D-BSSE/UniBasel facility under standard chow, water, enrichment, and light/dark cycles.

To infer tissue boundaries and to mark the 3D cell morphology of the epithelium, the following transgenic alleles were used: Shh-cre (Mouse genome informatics: Shh^tm1(EGFP/cre)cjt)^ (Harfe et al., 2004), ROSA^mT/mG^ (mouse genome informatics: Gt(ROSA)26Sor^tm4^(^ACTB-tdTomato,-EGFP)Luo^)(Muzumdar et al., 2007) and Hoxb7/myr-Venus (mouse genome informatics: Tg(Hoxb7-Venus^*^)17Cos) (Chi et al., 2009).

The Shh-cre allele was used to drive Cre recombinase-mediated recombination of the ROSA^mT/mG^ allele. As recombined EGFP localizes to the cell membrane, and *Shh* is only expressed in the lung bud epithelium, individual cell morphology could be segmented. To generate a Shh-cre; ROSA^mT/mG^ double-transgenic line, Shh-cre mice heterozygous for the Shh^tm1(EGFP/cre)Cjt)^ allele were crossed with homozygous ROSA^mT/mG^ mice. F1 offspring were crossed again to obtain animals that are homozygous for the ROSA^mT/mG^ and heterozygous for the Shh-cre allele *(Shh^cre/+^; ROSA^mT/mG^)*. For the experiments in this study, *Shh^cre/+^; ROSA^mT/mG^* males were either crossed with RjOrl:SWISS wild type or ROSA^mT/mG^ females.

To obtain *Shh* knockout embryos, timed matings of *Shh^cre/+^; ROSA^mT/mG^* mice were set. A quarter of the collected embryos were homozygous for the Shh-cre allele and therefore did not carry the wild type *Shh* gene. Shh knockout embryos displayed early defects in vertebrate embryonic tissues, including the brain, limbs as well as in lung patterning and branching.

For the kidney explant cultures, homozygous Hoxb7/myr-Venus males were crossed with RjOrl:SWISS wild type females because of higher pregnancy rates and larger litters.

### 2D organ culture and time-lapse epifluorescence imaging

Pregnant female mice were sacrificed according to institutional guidelines at the desired embryonic stages. All embryos were sorted using a fluorescence stereomicroscope (Leica Microsystems) according to the desired fluorescent reporters. The lungs and metanephric kidneys were dissected in ice-cold PBS and cultured at the liquid-air interface (Grobstein, 1956) using a trans-well membrane culture system supplemented with explant culture medium (DMEM/F12, 10% FBS (Sigma-Aldrich Chemie GmbH; F9665-500ML), 1x GlutaMAX (Life Technologies Europe BV; A1286001), 1x Penicillin-Streptomycin (Roche; 11074440001)). For the FGFR inhibitor lung cultures, the culture medium was supplemented with 3 μM or 5 μM SU5402 (Sigma-Aldrich, SML0443) before live imaging. For the lung cultures without a trachea, the trachea was cut right below the larynx or above the carina before live imaging. For the MMP inhibitor lung cultures, the culture medium was supplemented with 2.5 μM or 20 μM GM6001 (Abcam; ab120845) before live imaging.

To investigate the developmental dynamics of isolated lung bud and ureteric bud (UB) epithelia, mesenchymal tissue was enzymatically removed (Fig. 3D). The experimental procedure was adapted from (Nogawa and Ito, 1995; Yuri et al., 2017). Dissected lungs and kidneys were washed in DMEM/F12 and then incubated in 200ng/ml Collagenase/Dispase (Roche; 10269638001) in DMEM/F12 for 10 min (kidney) or 12 min (lung) in an incubator (37 °C 5% CO2), washed once in DMEM/F12 with 50% FBS and then placed in 100% FBS on ice for 10min to stop the enzymatic reaction. The mesenchyme was removed using tungsten needles. Isolated UBs were embedded in 50% Growth-Factor Reduced Matrigel (GFRMG) (Corning; 354230) and 50% explant culture medium supplemented with 4 ng/ml rhGDNF (R&D; 212-GD), 200 ng/ml human FGF-acidic (Peprotech; 100-17A), 200 ng/ml murine R-Spondin (Peprotech; 315-32) and 0.2 μM retinoic acid (Sigma-Aldrich; R2625). A 20 μl drop of this mixture was placed on a trans-well filter, and an isolated UB was carefully moved into the drop, with as little medium as possible, and allowed to sink atop of the membrane to largely restrict growth to two dimensions. Supplemented explant culture medium without GFRMG was added to the well, and the plate was placed in an incubator (37 °C, 5% CO2) for at least 2h prior to live imaging. For the pERK staining (Fig. 3), UBs were isolated and cultured as described, however, the filter membrane was omitted and the samples embedded in a hanging drop of 1:1 GFRMG/explant culture medium + growth factors. Isolated lung buds were placed in a 20 μl drop of GFRMG in a tissue-culture treated plate (Corning; 3513) solidified upside down (hanging-drop) in an incubator (37 °C, 5% CO2) for 20min, then another drop of GFRMG (15 μl) was added on top. After 30-45min, this drop was overlaid with explant culture medium supplemented with 100 ng/ml human FGF10 (Peprotech; 100-26), 50ng/ml human FGF-acidic, 50 ng/ml R-Spondin and 2 μM CHIR99021 (Sigma-Aldrich; SML1046).

Automated time-lapse imaging of the cultured explants was performed using a Nikon epifluorescence microscope, which was equipped with an environmental control chamber. A 10x/0.3na Plan Fluor Ph1 DLL objective with 16 mm working distance (Nikon; MRH10101) and a 475/28 excitation filter was used to capture GFP or myr-Venus fluorescence in the epithelium, and a 549/15 excitation filter was used to capture RFP fluorescence in the mesenchyme. Images were acquired using a Hamamatsu Orca Flash 4.0 V2 camera at set positions every hour for 48h (SU5402 and trachea cultures) or 60h (all other cultures with medium changes after 48h).

### Quantification of fluid flow in the lung lumen

Injection needles were pulled from borosilicate thin wall capillaries (Warner Instruments; 300038) using a microneedle puller (Narishige Group; PN-30). E11-11.5 lungs were placed on a microscopy slide mounted in a Leitz Labovert inverted stereo microscope. FluoSpheres polystyrene microspheres (1μm, yellow-green fluorescent (505/515)) (Invitrogen; F13081) were diluted 1:100 in sterile 1x DPBS (Life Technologies; 14040174), and a Leitz micromanipulator connected to an Eppendorf FemtoJet 4i (Eppendorf; 5252000013) was used to inject a small volume of the bead suspension into the intraluminal space of the most distal tip of the left or right bronchus. We used a low-volume organ culture technique to minimize the distance between objective and sample to improve image quality and reduce phototoxic effects on the sample. Briefly, samples were washed in explant culture medium, placed in a Falcon center-well organ culture dish (Corning; 353037) in 120μl pre-warmed explant culture medium. 60μl of a 1:1 mix of medium and GFRMG was added around the samples for support. Samples were allowed to rest in the incubator for at least 15min before transferring them to a Nikon Eclipse Ti2 inverted microscope equipped with a Nikon Crest X-light V3 spinning disk confocal system, a Photometrics Prime 95B 25mm sCMOS camera and an environmental control chamber set to 37 °C, 5% CO2. Images were acquired using a 20x/0.75na CFI Plan Apo Lambda objective (Nikon Instruments; MRD00205). To image the fluorescent beads, a 80μm Z-stack (10μm Z-steps) and a brightfield image of the lung at the centre position were acquired at each time step. Both ‘no delay’ imaging with around 7 second intervals for a duration of 10min, or 1-2h acquisitions at 1 min time steps were performed). For the measurement of the fluid flow, we used time lapses acquired at 1 min steps to reduce adverse effects of constant light exposure on the sample. The NIS-Elements (version 5.21.00) ‘Focused Image’ feature (Z-Map method: Balanced; Zero-based) was used to create a Z-projection of the bead channel. The beads were tracked using Imaris and Imaris Track (Bitplane, South Windsor, CT, USA) by creating a spot object and tracking the segmented spots over time using Imaris’ ‘autoregressive motion’ algorithm. The Imaris MeasurementPro Statistics feature was used to export the maximum track speed.

### Immunofluorescence, optical clearing and light-sheet imaging

To detect phosphorylated MAP Kinase Erk1 and Erk2 (pERK) (Fig.3) in E12.5 lungs and kidneys and in mesenchyme-free E11.5 UBs cultured for 72h, immunostaining was performed as described in (Menshykau et al., 2019). Mesenchyme-free UB cultures were treated with Corning Cell Recovery Solution (Corning; 354253) for 10min at 4 °C to digest the Matrigel prior to fixation. After staining, an additional post-fixation step with 4% paraformaldehyde (PFA) for 20min at 4 °C was performed prior to optical clearing to preserve fluorescence. For the actin and fibronectin immunostaining (Fig. 3F,G, Fig. 5B,C), lung samples were preblocked (1% BSA, 10% donkey serum (Sigma-Aldrich; D9663), 0.5% Triton-X (Sigma-Aldrich; T8787) in 1x PBS) overnight, incubated with primary antibody diluted 1:200 (Anti-Actin: Abcam; ab179467) or 1:150 (Anti-Fibronectin: Abcam; ab2413) in blocking solution for two days, washed with PBS during one day, incubated with a fluorescently conjugated secondary antibody diluted 1:500 overnight (Alexa Fluor 555: Invitrogen; A-31572) or for two days (Alexa Fluor Plus 647: Invitrogen; A-32795), and washed with PBS during one day. Samples were post-fixed in 4% PFA for 20min and washed in PBS.

Whole-mount tissue clearing of dissected embryonic explants was performed with the Clear Unobstructed Brain/Body Imaging Cocktails and Computational Analysis (CUBIC) protocol as specified by the authors (Susaki et al., 2015). Clearing times in reagents for decolouring, delipidation, permeation (CUBIC-1), and refractive index (RI) matching (CUBIC-2) were adjusted to maximize clearing efficiency and minimize quenching. Following a 1h fixation in 4% PFA, samples were incubated in 1/2 CUBIC-1 (CUBIC-1:H_2_O=1:1) for two-four days, and in 1X CUBIC-1 until they became transparent. All samples were subsequently washed several times in PBS. The actin immunostaining was performed after the CUBIC-1 clearing steps. All samples were then treated with 1/2 CUBIC-2 (CUBIC-2:PBS=1:1) for two-four days. Lastly, incubation in 1X CUBIC-2 was done until the desired transparency was achieved. CUBIC-1 steps were performed on a shaker at 37 °C while CUBIC-2 steps at RT. Cleared samples were embedded in 2% low-melting-point solid agarose cylinders, and immersed in CUBIC-2 overnight. 3D image stacks were acquired using a Zeiss Lightsheet Z.1 SPIM microscope using a 20x/1.0 clearing objective.

### 3D organ culture, mounting, and time-lapse light-sheet imaging

Following dissection in DPBS at RT, all lung explants were cultured in sterile Dulbecco’s modified Eagle’s medium w/o phenol red (DMEM) (Life Technologies Europe BV; 11039021) containing 10% Fetal Bovine Serum (FBS) (Sigma-Aldrich Chemie GmbH; F9665-500ML), 1% Glutamax (Life Technologies Europe BV; A1286001), and 1% penicillin/streptomycin (Life Technologies Europe BV; 10378-016). All specimens were equilibrated at 37 °C with 5% CO_2_ in a humidified incubator for 1h.

In the meantime, LMP hollow agarose cylinders were prepared according to (Udan et al., 2014). Hollow cylinders allow for unencumbered 3D embryonic growth, minimize tissue drift, enable imaging from multiple orientations, and allow for better nutrients and gas perfusion. Within a hollow cylinder, a single specimen was suspended in undiluted Matrigel (VWR International GmbH; 734-1101) to recapitulate the in-vivo microenvironment. All cylinders were kept at 37°C with 5% CO_2_ in culture media for 1h prior to mounting.

For an overnight culture, the imaging chamber was prepared first by sonication at 80 °C and subsequent washes in ethanol and sterile PBS. After the chamber was assembled, culture medium and allowed to equilibrate at 37 °C with 5% CO_2_ for at least 2h before a cylinder was mounted for imaging. Furthermore, to compensate for evaporation over time and maintain a fresh culture media environment, peristaltic pumps were installed to supply 0.4 ml and extract 0.2 ml of culture medium per hour. Each lung explant was then aligned with the focal plane within the centre of a thin light-sheet to enable fine optical sectioning with optimal lateral resolution. For this study, all samples were imaged using a 20x/1.0 Plan-APO water immersion objective.

### Segmentation of explant morphology in 3D

Light-sheet datasets were transferred to a remote storage server and processed in a remote workstation (Intel Xeon CPU E5-2650 with 512 GB memory). Deconvolution via Huygens Professional software (SVI) improved overall contrast and resolution while Fiji (ImageJ v1.52t) (Schindelin et al., 2012) was used for accentuating cell membranes, enhancing local contrast, and removing background fluorescence. To extract 3D morphological measurements, the length was measured along the centre of Imaris 9.1.2 (Bitplane, South Windsor, CT, USA) iso-surfaces, and cross-sections of tubular bronchial portions were masked and exported into Fiji, where 2D circumference was calculated and averaged over the tube.

### Segmentation and skeletonization of 2D culture datasets

Epifluorescence images of embryonic lung and kidney explants were processed in Fiji (ImageJ v1.52t) (Schindelin et al., 2012). Before segmentation, local image contrast was increased, and image background subtracted. Images were then binarized using a global thresholding method, and boundaries were smoothened by applying a gaussian blur filter. Skeletonization of binarized images was performed with the Skeletonize3D plugin, the coordinates and length of all branches from a given skeleton were inferred with the AnalyzeSkeleton plugin (Arganda-Carreras et al., 2010), and branch widths were measured with the BoneJ plugin (v1.4.2) (Doube et al., 2010) (Supplementary Fig. S8).

### Branch analysis software

Analysing the branching behaviour over time required the generation of branch lineages from a skeletonized time series. To this end, a MATLAB script was created to quantify branch length and width and track all branches over time (see https://gitlab.ethz.ch/chrlang/branching-analysis).

### Continuum-mechanical simulations of epithelial tube collapse

A full technical description of the custom finite element simulation framework that we employed to simulate epithelial tube collapse (Fig. 5) can be found in (Vetter et al., 2013). Originally developed for more general linearly (visco-)elastic thin structures, it allows for efficient transient simulation of thickness-preserving epithelial tissues in 2D. Since the full set of equations solved fills pages, we repeat only the most important aspects here. We minimized the total two-dimensional elastic energy *U* of the tissue cross section, given by

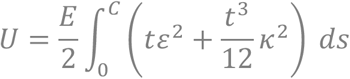

In this formalism, the tissue thickness *t* is a constant parameter that does not change as the tissue deforms. Young’s modulus *E* sets the energy scale in the simulations, affecting all deformation modes equally. *C = 2πR* denotes the total tubular circumference as measured along the tissue midline (Fig. 5A), *ds* the infinitesimal arclength element, *ε* the hoop Cauchy strain, and *κ* the tissue curvature on the midline. The shape of the epithelium was represented by third-order finite beam elements using cubic Hermite splines, which were embedded in a corotational formulation to allow for large (geometrically nonlinear) deformations. In addition to *U*, the employed third-order beams contain an energy term for transverse shear that depends on the second modulus of isotropic linear elasticity, Poisson’s ratio *ν* (see Vetter et al., 2013 for details). A second-order Newmark method in predictor-corrector form was used to integrate Newton’s second law of motion,

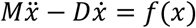

Here, *M* denotes the finite element mass matrix, *D* the viscous damping matrix, *x* the vector holding the translational and rotational degrees of freedom, and *f* the total generalized force vector including all forces from elasticity (–∇_x_U), contact, applied pressure, and the volume constraint. Elastic contact between tube segments and between the epithelium and the rigid clamps was modelled using Hertzian contact mechanics where applicable, as also detailed in (Vetter et al., 2013). In the volume-controlled collapse scenario, the volume enclosed by the epithelium was imposed with a Lagrange multiplier and reduced linearly over time. In the pressure-controlled simulation, a net pressure difference *ΔP* was applied and slowly increased linearly over time until the critical buckling threshold was surpassed and the tube collapsed. In the simulation with clamps, parallel rigid walls were approaching one another at a low constant speed until they were 2.5*t* apart. To build the mass matrix *M*, we assumed a homogeneous mass density ρ in the tissue. The damping matrix was set to *D = M/τ* where τ sets the viscous relaxation time of the tissue. The equations of motion were solved until static equilibrium was reached *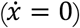*, yielding the configurations shown in Fig. 5. Each simulation used a spatial resolution of 100 finite elements along the circumference. All relevant simulation parameters are listed in Supplementary Table 1.

### Estimation of shear stress levels in embryonic lungs

The wall shear stress on the apical side of a closed lung tube was estimated with the finite element software Comsol Multiphysics (v5.4). As a reference geometry, we used a section of a lung tube originating from an E11.5 embryo of the Shh^cre/+^ line described above. We chose a 400 μm segment from the carina to the first branch of the left lung lobe (LL1). A surface encompassing the lumen of this tube was extracted from the light-sheet images using the ‘automatic surface creation’ feature of Imaris. The intense fluorescence in this region imposed us to include, in addition to the apical surface, a part of the tissue surrounding the lumen. Consequently, the geometry obtained had a cross-sectional opening of about 20 μm along its minor axis. To reduce the luminal width back to the measured range, the geometry was scaled down along the Z-axis in Paraview (v5.7.0). Different scaling factors were employed to obtain several versions of this geometry with varying degrees of lumen opening. Fluid flow inside the lumen was simulated by solving the Stokes equations for incompressible Newtonian fluids. The dynamic viscosity of the fluid was set to 0.016 Pa·s (Bokka et al., 2015), and its mass density was set to that of water (1000 kg/m^3^). A no-slip condition was assumed at the interface between the fluid and the surface of the epithelium. To generate fluid motion, the average flow velocity at the inlet was set to 0.364 μm/s as measured in the lungs, and the pressure at the outlet was maintained at 1 atm. The average wall shear stress value on the apical surface was measured with a boundary probe. We repeated the computations with the previously estimated volume flow rate of 420 μm^3^/s (George et al., 2015).

### Cell-based simulations of shear stress effect

We simulated the growth of the lung epithelium subjected to shear stress using a vertex model available in the Chaste framework (Fletcher et al., 2013; Mirams et al., 2013). The dynamics of the vertices were derived from the potential energy of the system, as previously proposed (Nagai and Honda, 2009). The parameters used to calibrate the model are given in Supplementary Table 2. The initial tissue configuration was comprised of 100 cells arranged in a honeycomb with an aspect ratio of 1. While the cells located in between were proliferative, the cells on the top-most and bottom-most layers were differentiated and never underwent mitosis. To simulate the elongating effect of shear stress, external forces were applied to the vertices of the differentiated cells. Since the movement of luminal fluid is unidirectional in lung tubes, external forces were applied in the direction of outgrowth. To generate the elongating effect of shear stress vertical upward forces were applied to the differentiated cells of the top layer, while vertical downward forces were applied to the cells of the bottom layer. The magnitudes of these forces ranged from 0 to 2 a.u. as specified in figure 7. The tissue boundary was left free to move without constraints. We employed MorphoGraphX to 2.5D segment the apical layer of E11.5 lungs, and extract both the area and number of neighbour distributions (Barbier de Reuille et al., 2015). We observed an apical area coefficient of variation (CV) of 0.6 and a percentage of hexagons of 30%. As these two parameters are dependent on cell growth and division rates (Kokic et al., 2019; Sahlin and Jonsson, 2010), the cell division areas and cell cycle durations were drawn randomly from normal distributions to reproduce equivalent statistics *in-silico* (see Supplementary Table 2). Furthermore, in compliance with Hertwig’s rule (Hertwig 1884), the cell division axis at mitosis was set to be the shortest axis through the cell centroid. All simulation durations were set to 48.0 a.u.

## Code Availability

The source code for the cell-based simulations and the model file for the fluid flow simulations are freely available as a git repository at https://git.bsse.ethz.ch/iber/Publications/2021_conrad_runser_biased-elongation.git.

## Lot numbers

- Anti-Actin antibody: GR206286-25
- Anti-Fibronectin antibody: GR3323518-3
- Phospho-p44/42 MAPK (Erk1/2) (Thr202/Tyr204) antibody: 15
- SU-5402: 104M4713V
- Recombinant human FGF-acidic: 031207 K1014
- Recombinant human FGF10: 0518162-1 K0518
- Recombinant murine R-Spondin: 0516620 E0516
- Recombinant human GDNF: VQ2517081
- Collagenase/Dispase: 11488501
- GM6001: APN17122-1-1

## AUTHOR CONTRIBUTIONS

DI conceived the study. HG obtained the light-sheet microscopy data in Figures 1 and 4, and, together with OM, the pERK staining in Figure 3B,C. MD obtained the 3D live imaging data in Figure 4, with support from OM and HG. LC obtained and analysed the lung and kidney culture data in Figures 1 and 2, generated the GM6001 culture data in Figure 3, the staining in Figures 3D,F,G and 5, with support from LK, and the fluid flow measurements in Figure 6. CL obtained and analysed the lung cultures in Figures 1, 3 and 6. LC and CL wrote a Fiji script for the image processing of the filter cultures. MD and CL wrote a MATLAB script for the 2D analysis of the filter cultures. AS obtained, processed, and analysed the lightsheet microscopy data for Figures 6 and 7. SR developed the Comsol and Chaste simulations, with support from RV. RV developed the continuum mechanics simulations. DI wrote the manuscript, with contributions from RV; HG, SR, LC, CL, RV wrote the Methods section and figure legends; all authors approved the final manuscript.

## ACKNOWLEGEMENTS

This work has been supported through an SNF Sinergia grant to DI. We thank Anna Stopka for preliminary quantifications of the biased lung tube elongation, and Alay Shah for segmenting the basal sides of the lung epithelium in MorphoGraphX. We greatly thank Tom Lummen (D-BSSE Single Cell Facility) for setting up the spinning disk confocal acquisition, and we are grateful to Makiko Seimiya for providing the micromanipulator system. We thank Jamie Davies and May Sallam for advice on the mesenchyme-free kidney culture, especially regarding the growth-factor concentrations.

## SUPPLEMENTARY TABLES

**Supplementary Table 1:**
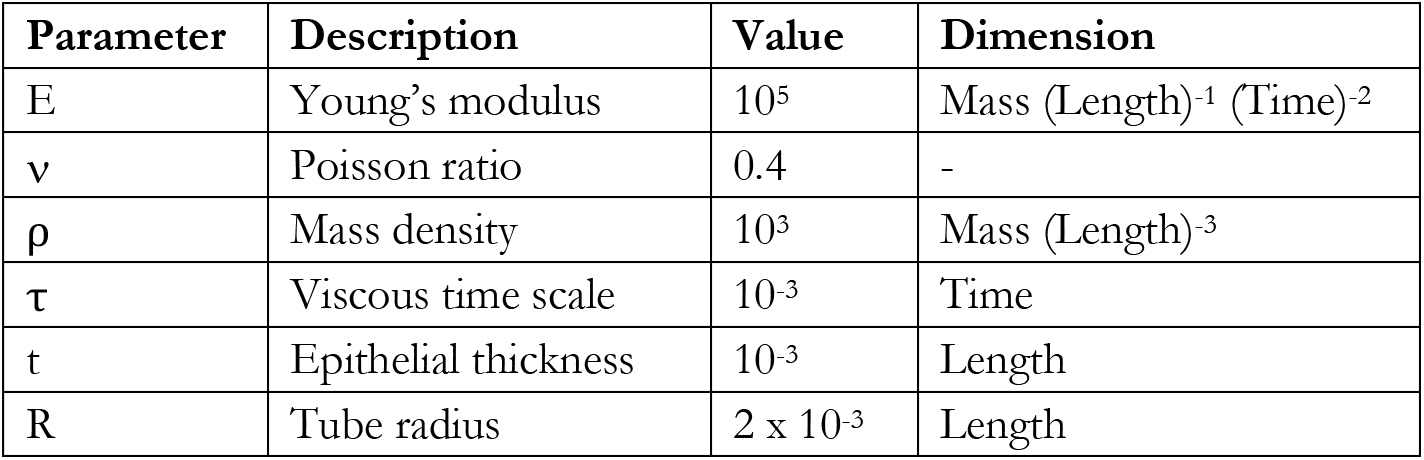
Parameter values used in the tube collapse simulations.

| Parameter | Description | Value | Dimension |
| --- | --- | --- | --- |
| E | Young's modulus | $10^5$ | Mass (Length)<sup>-1</sup> (Time)<sup>-2</sup> |
| $\nu$ | Poisson ratio | 0.4 | - |
| $\rho$ | Mass density | $10^3$ | Mass (Length)<sup>-3</sup> |
| $\tau$ | Viscous time scale | $10^{-3}$ | Time |
| t | Epithelial thickness | $10^{-3}$ | Length |
| R | Tube radius | $2 \times 10^{-3}$ | Length |

**Supplementary Table 2:**
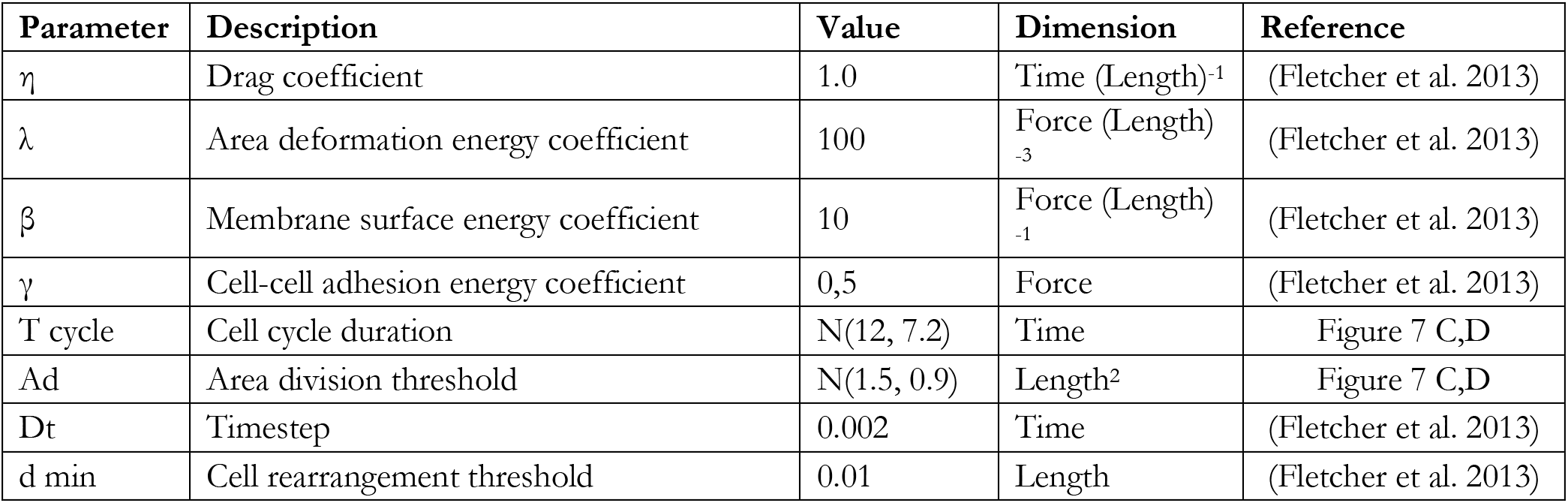
Parameter values used in the Chaste simulations.

| Parameter | Description | Value | Dimension | Reference |
| --- | --- | --- | --- | --- |
| $\eta$ | Drag coefficient | 1.0 | Time (Length)<sup>-1</sup> | (Fletcher et al. 2013) |
| $\lambda$ | Area deformation energy coefficient | 100 | Force (Length)<sup>-3</sup> | (Fletcher et al. 2013) |
| $\beta$ | Membrane surface energy coefficient | 10 | Force (Length)<sup>-1</sup> | (Fletcher et al. 2013) |
| $\gamma$ | Cell-cell adhesion energy coefficient | 0,5 | Force | (Fletcher et al. 2013) |
| T cycle | Cell cycle duration | N(12, 7.2) | Time | Figure 7 C,D |
| Ad | Area division threshold | N(1.5, 0.9) | Length<sup>2</sup> | Figure 7 C,D |
| Dt | Timestep | 0.002 | Time | (Fletcher et al. 2013) |
| d min | Cell rearrangement threshold | 0.01 | Length | (Fletcher et al. 2013) |

## SUPPLEMENTARY FIGURES

**Supplementary Figure 1:**
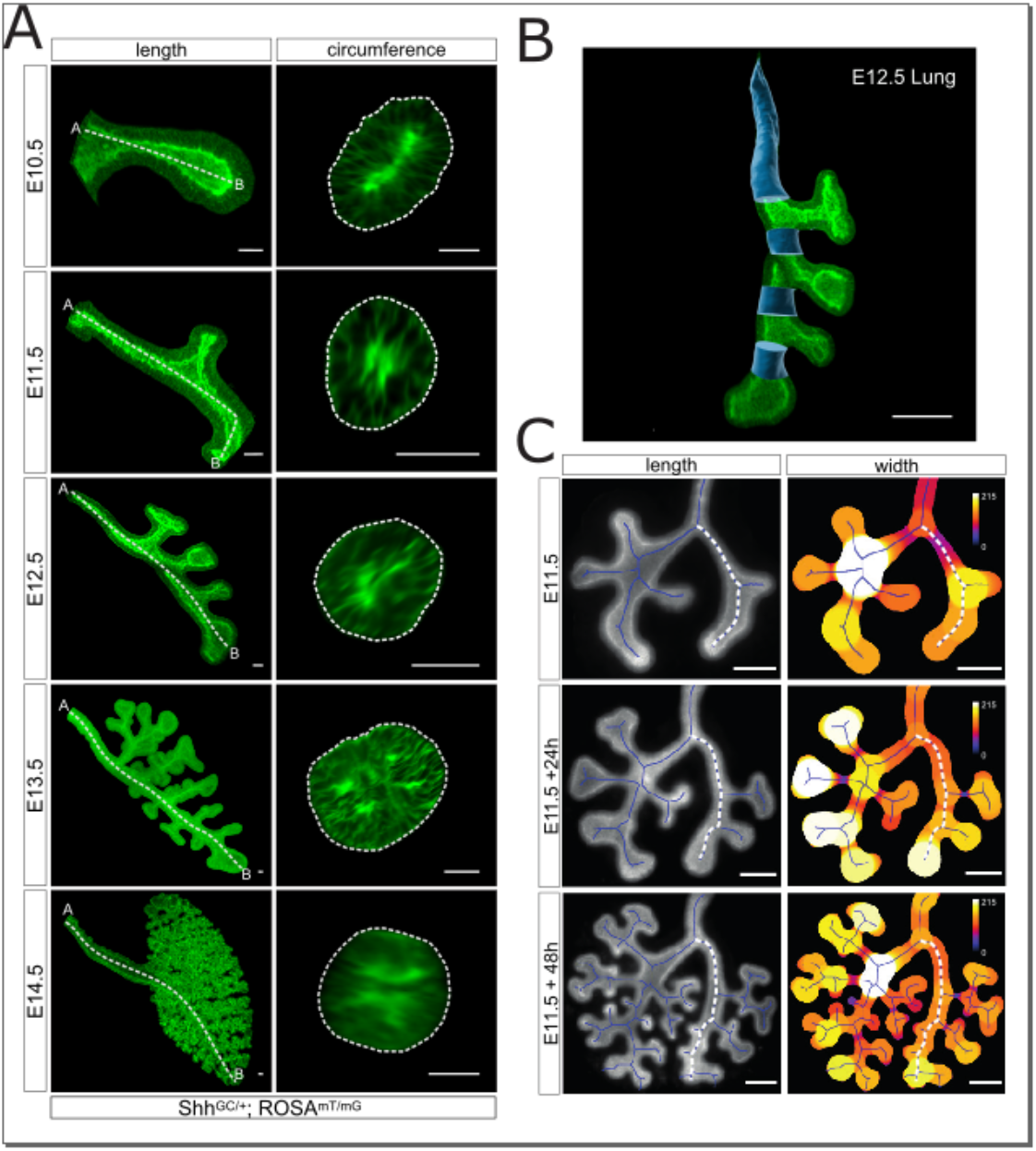
Measuring epithelial morphology in growing epithelial tubes. **(A)** 3D morphometric measurements of branch length and average circumference for a developmental timeline of a mouse lung. Specimens were serially isolated between E10.5 and E14.5 and carried the Shh^GC/+^; ROSA^mT/mG^ reporter, which enabled the visualization of the embryonic epithelium. Branch length was measured from below the carina to the most distal tip, while average branch circumference was calculated for tubular cross-sections. Scale bars 50 μm. **(B)** E12.5 mouse lung with iso-surface overlays denoting tubular sections used for morphometric quantifications (blue). Scale bar 200 μm. **(C)** 2D morphometric measurements of length and diameter for an E11.5 lung cultured on a filter for 48h. Width scale in μm. Scale bars 200 μm.

**Supplementary Figure 2:**
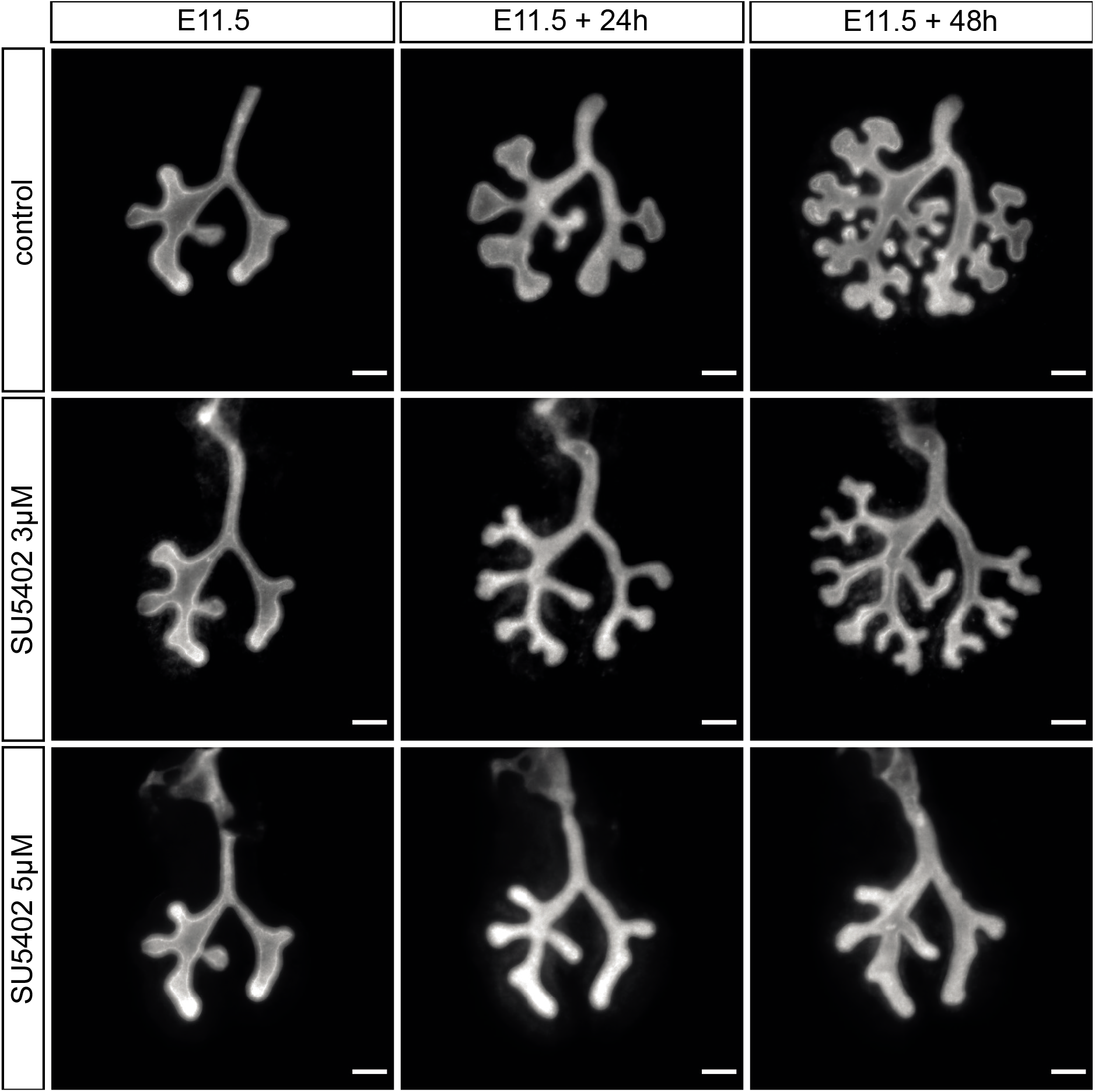
Lung explant cultures with FGFR inhibitor SU5402. Culture of E11.5 embryonic lungs under control conditions and under different concentrations of the FGFR inhibitor SU5402 for 48h. Scale bars 200 μm.

**Supplementary Figure 3:**
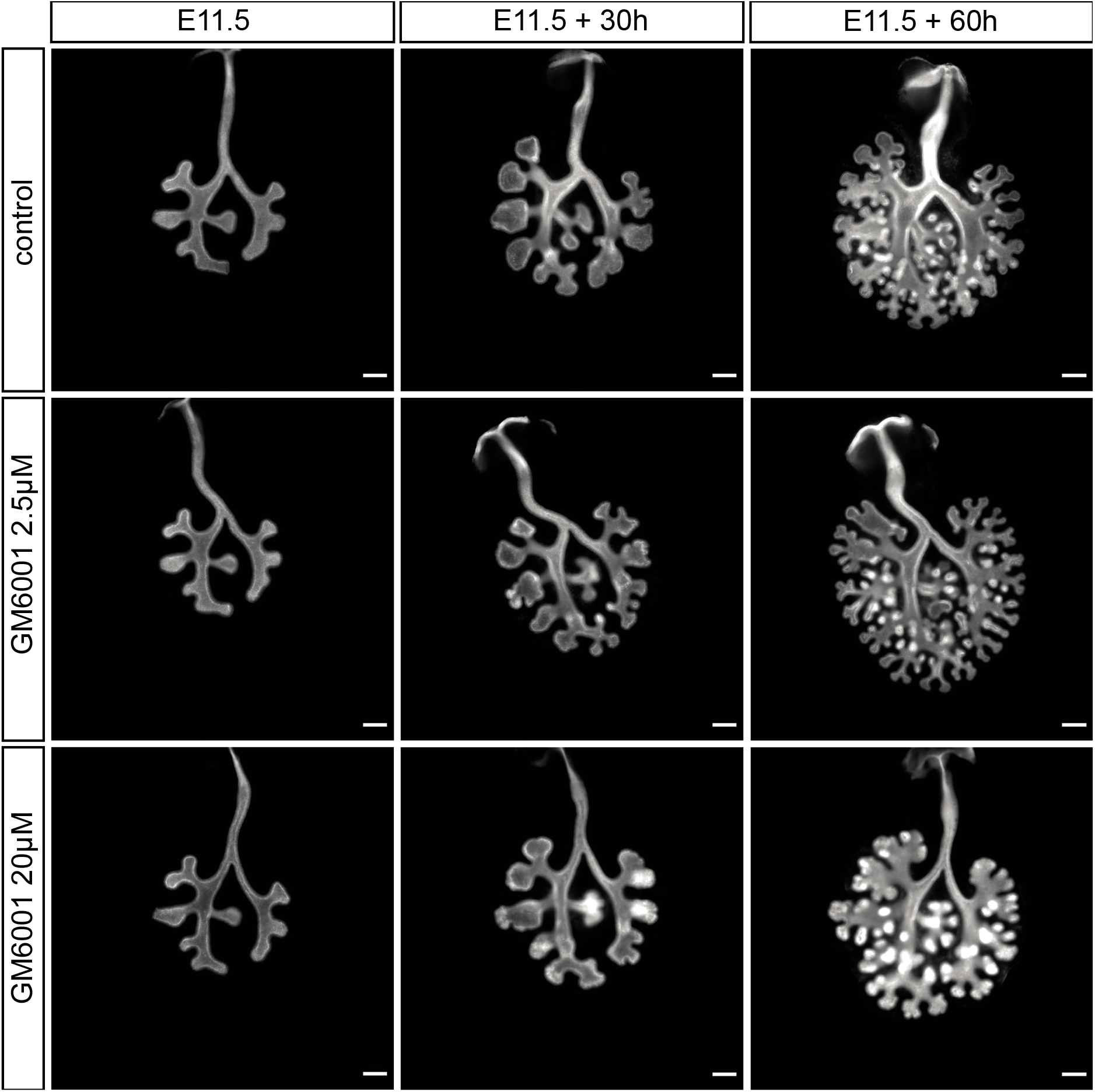
Lung explant cultures with MMP inhibitor GM6001. Culture of E11.5 embryonic lungs under control conditions and under different concentrations of the MMP inhibitor GM6001 for 60h. Scale bars 200 μm.

**Supplementary Figure 4.**
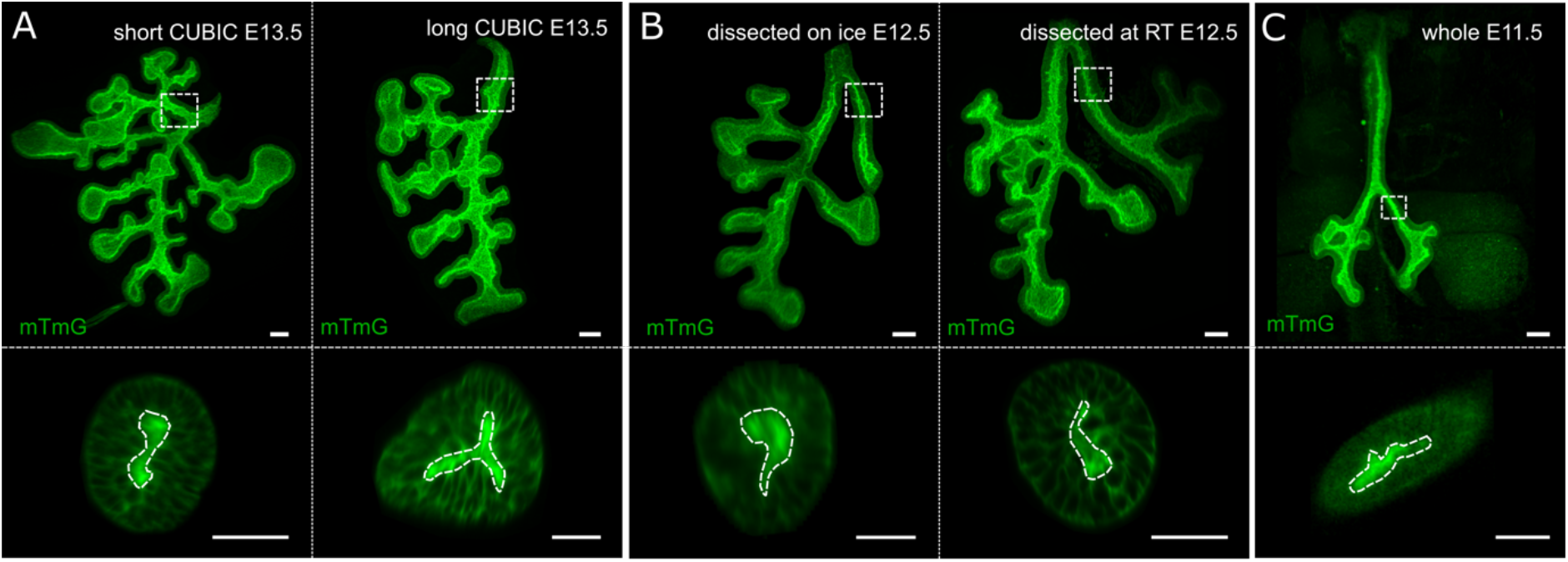
Narrow luminal spaces are not the result of dissection /clearing conditions. 3D rendering of mouse lung explants (top) and cross-sectional slices (bottom) of specimens **(A)** cleared using a short CUBIC regimen (2 days in reagent-1, 2 days in reagent-2) and a long CUBIC regimen (4 days in reagent-1, 4 days in reagent-2) displaying narrow luminal spaces. Similarly, **(B)** both explants dissected on ice and at room temperature (RT), and cleared using a short CUBIC regimen, also showed collapsed lumens. Furthermore, the same luminal morphology was observed in **(C)** CUBIC cleared whole-embryos imaged through thoracic cavity. Scale bars 100 μm.

**Supplementary Figure 5:**
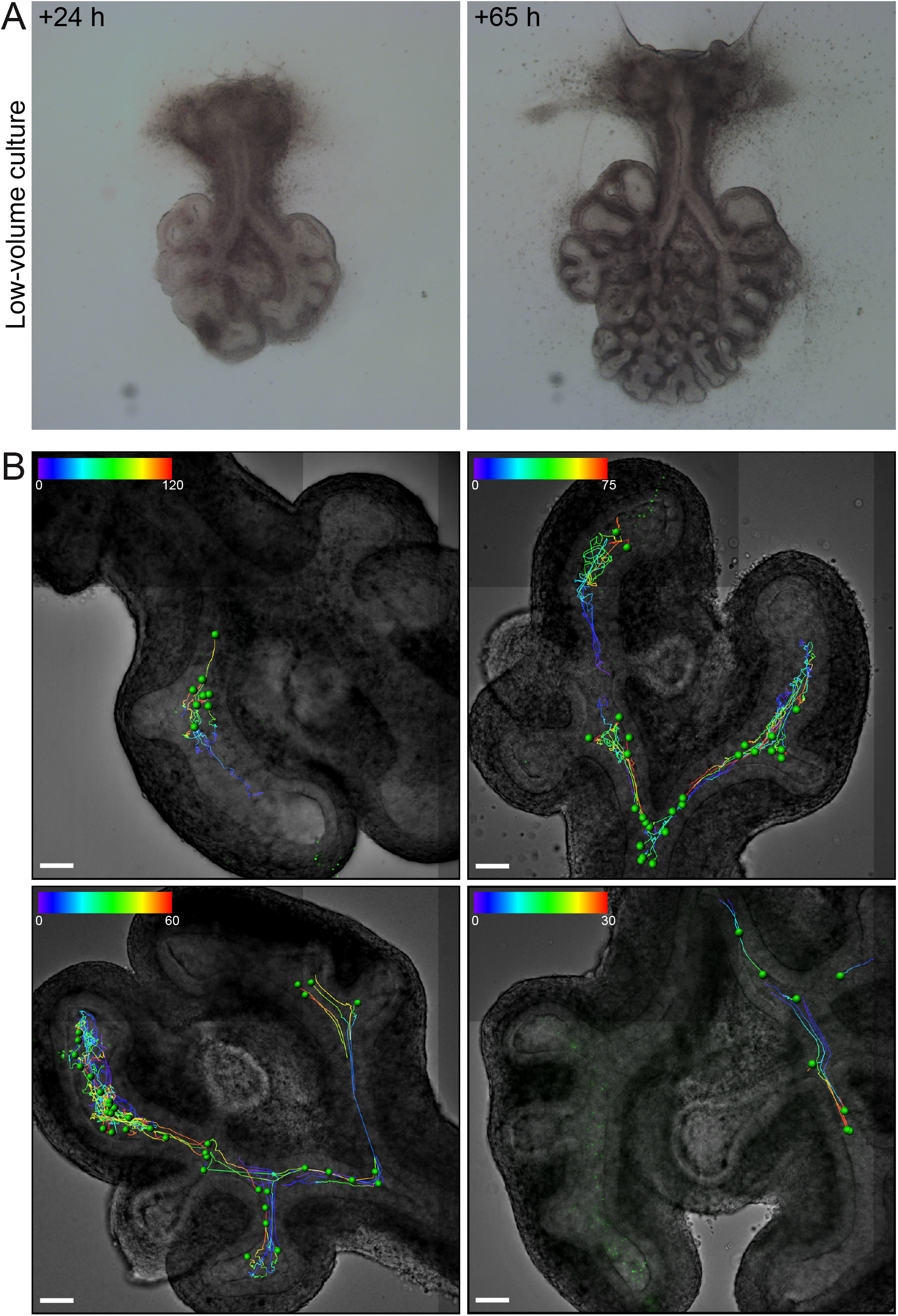
Bead time-lapse imaging and tracking in low-volume cultured E11.5 lungs. **(A)** Brightfield images of E11.5 lungs after injection and spinning disk confocal imaging after 24 h and 65 h continued low-volume culture. The culture medium contains 20% Matrigel. The liquid explant culture medium was replaced after spinning disk confocal imaging and after one day of culture. **(B)** Brightfield images of injected E11.5 lungs overlayed with the confocal bead images and the bead tracking (Imaris) of four experiments. Green dots mark the bead position at the end of the track. Colour bar denotes time (min) of tracks; scale bar 70 μm.

**Supplementary Figure 6:**
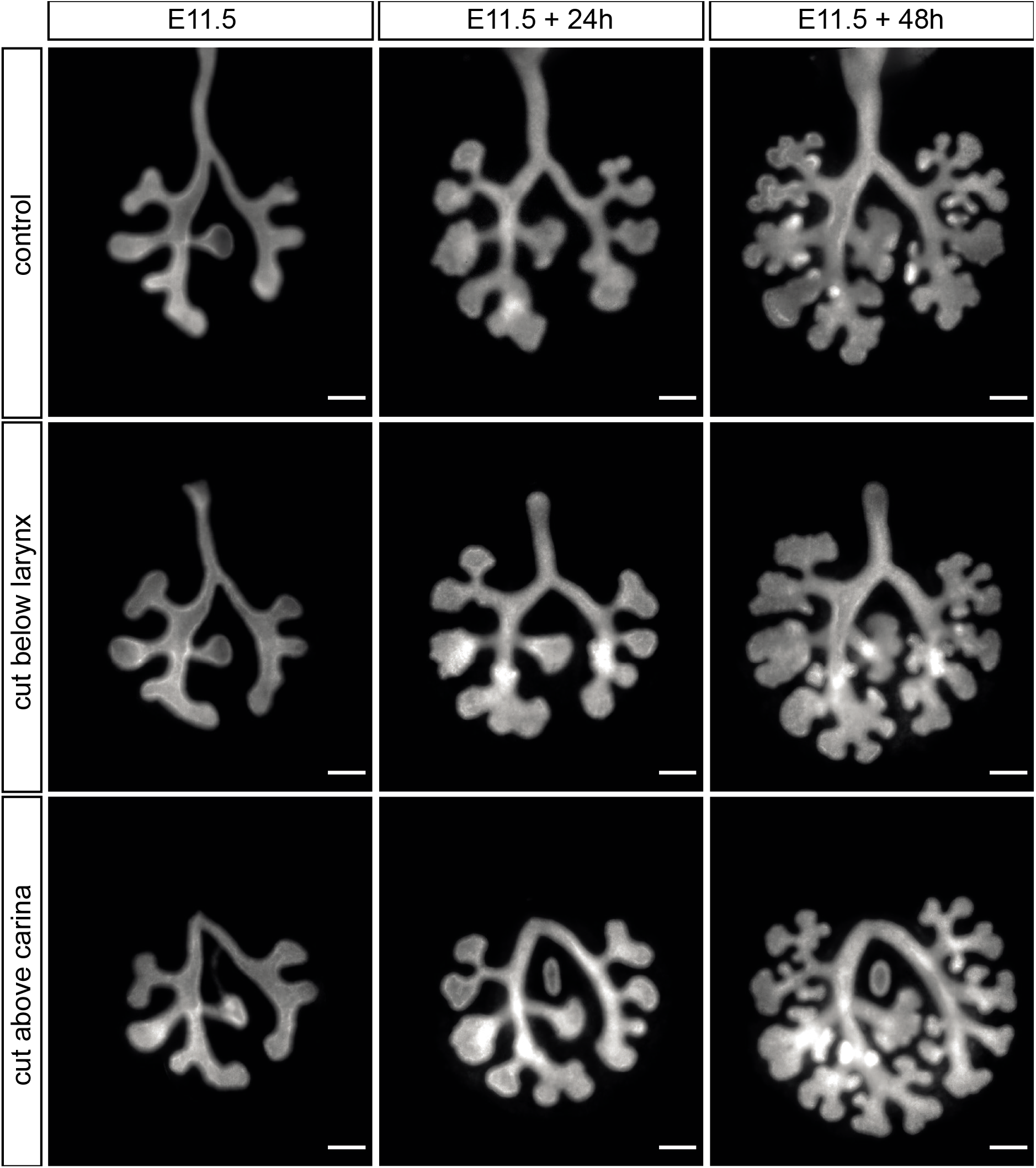
Lung explant cultures with intact and cut trachea. Culture of E11.5 embryonic lungs with intact trachea as control condition and with tracheas either cut below the larynx or above the carina for 48h. Altering the tracheal length does not impact on branching morphogenesis. Scale bars 200 μm.

**Supplementary Figure 7:**
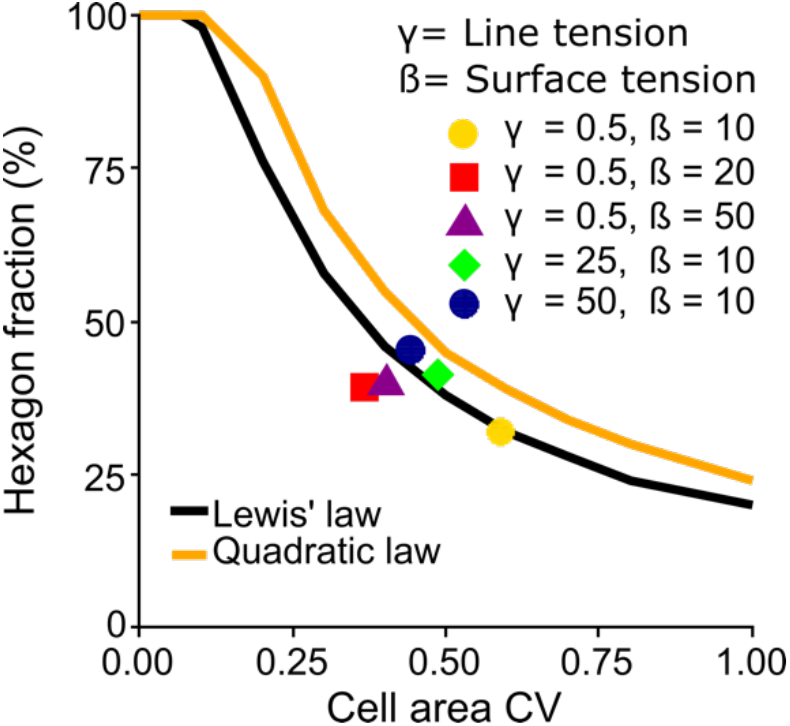
Impact of parameter variations in the Chaste simulations. Impact of the membrane surface energy β and the cell-cell adhesion energy g on the hexagon fraction and area CV in the Chaste simulations. As expected for a tissue representation of epithelia, the simulations reproduce the relationship between the hexagon fraction and the cell area CV as predicted based on Lewis’ law (black) or the quadratic law (yellow line) (Kokic et al., 2019).

**Supplementary Figure 8:**
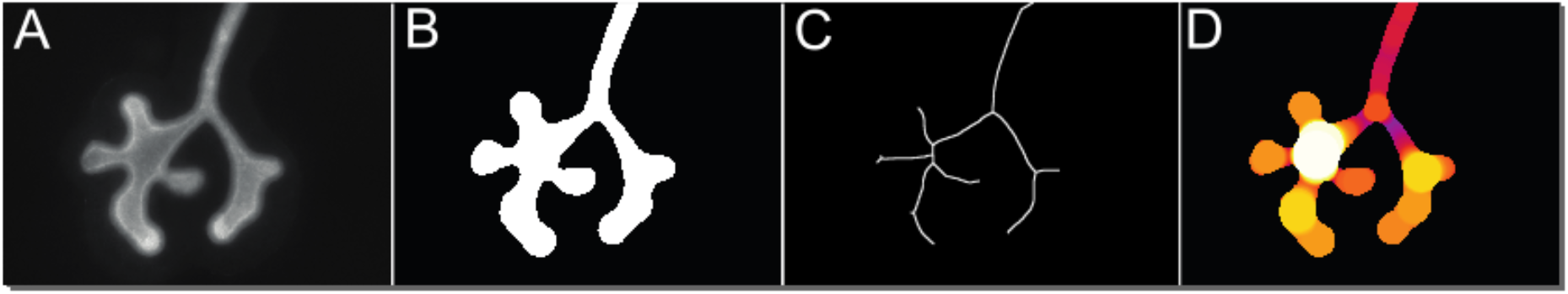
Image segmentation and skeletonization. Segmentation of raw images **(A)** resulted in binary images **(B),** which were then used to generate skeletons (**C**, inverted for illustration purposes). Thickness map images **(D)** were generated with the Fiji plugin BoneJ to infer branch widths.

## SUPPLEMENTARY VIDEOS

**Supplementary Video 1:**
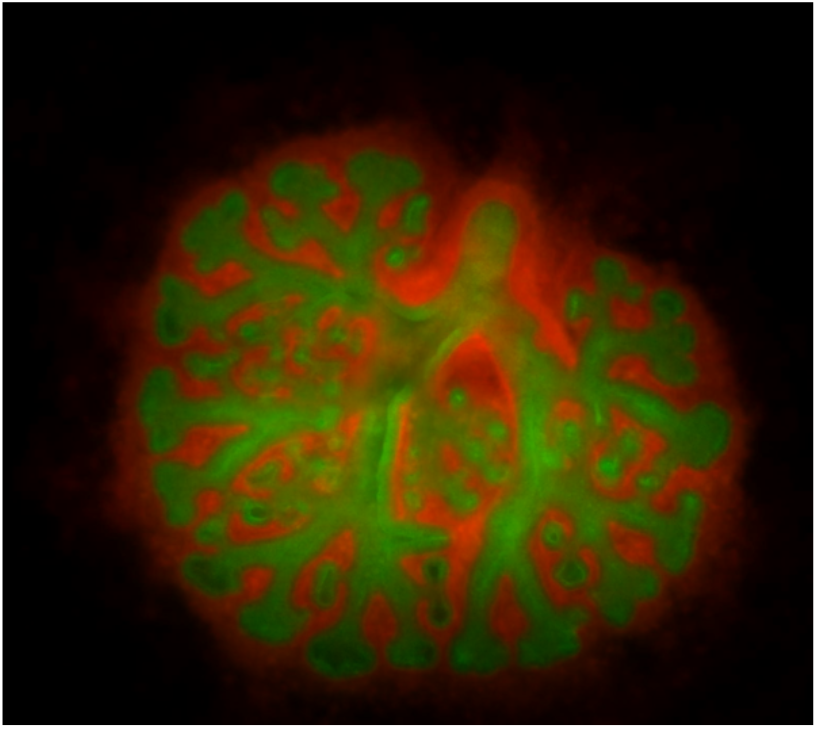
Lung explant culture. Time-lapse movie of a Shh^GC/+^; ROSA^mT/mG^ E11.5 lung cultured for 60h. Mesenchyme shown in red, *Shh-*expressing epithelium in green. (AVI 6.4 MB).

**Supplementary Video 2:**
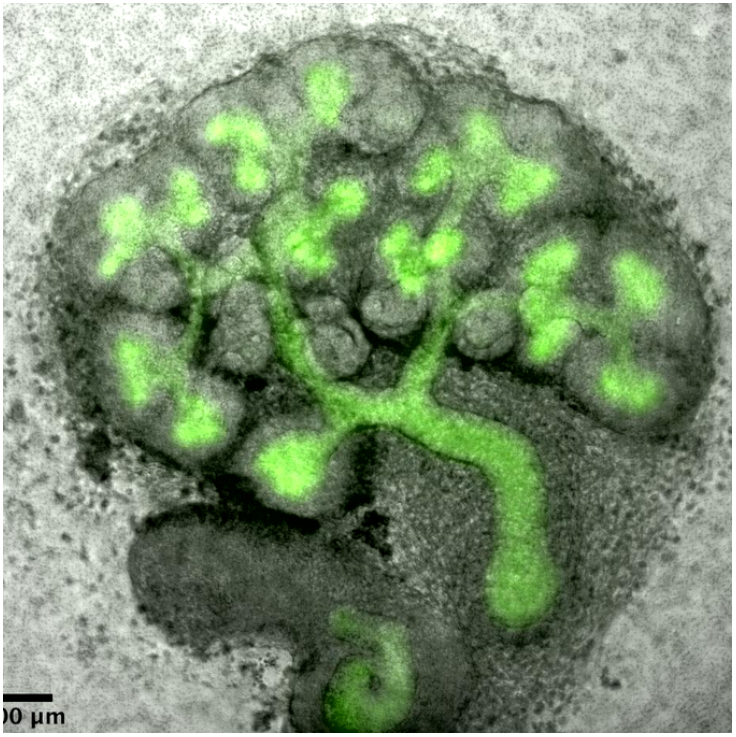
Kidney explant culture. Time-lapse movie of an E11.5 kidney cultured for 60h. HoxB7/myr-Venus expression is shown in green, brightfield in grey. (AVI 18.7 MB)

**Supplementary Video 3:**
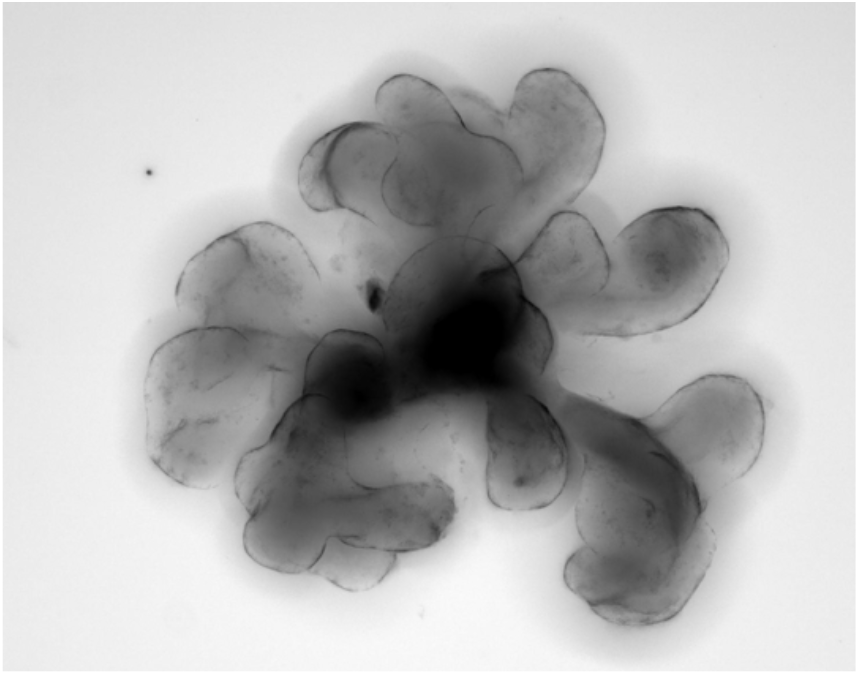
Mesenchyme-free lung bud culture. Time-lapse movie of a mesenchyme-free left lung bud dissected at E11.5 and cultured for 60h. (AVI 7.9 MB).

**Supplementary Video 4:**
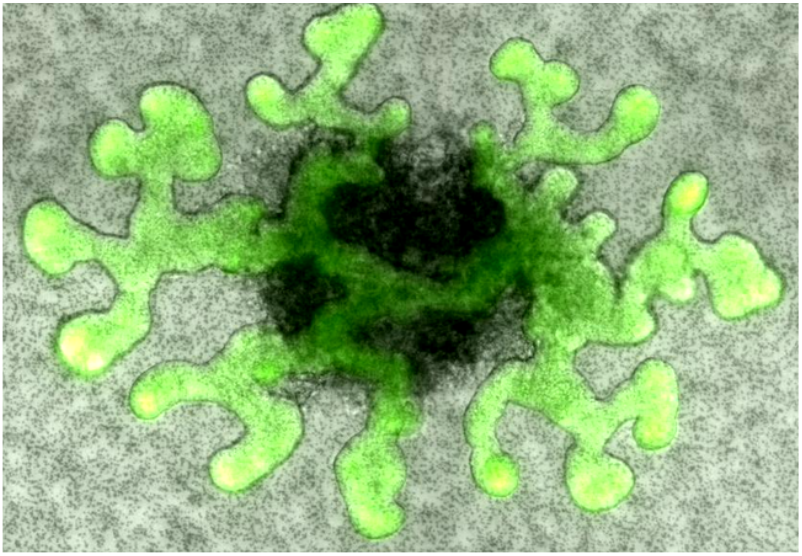
Mesenchyme-free ureteric bud culture. Time-lapse movie of a mesenchyme-free ureteric bud dissected at E11.5 and cultured for 60h. HoxB7/myr-Venus expression is shown in green, brightfield in grey. (AVI 10.8 MB).

**Supplementary Video 5:**
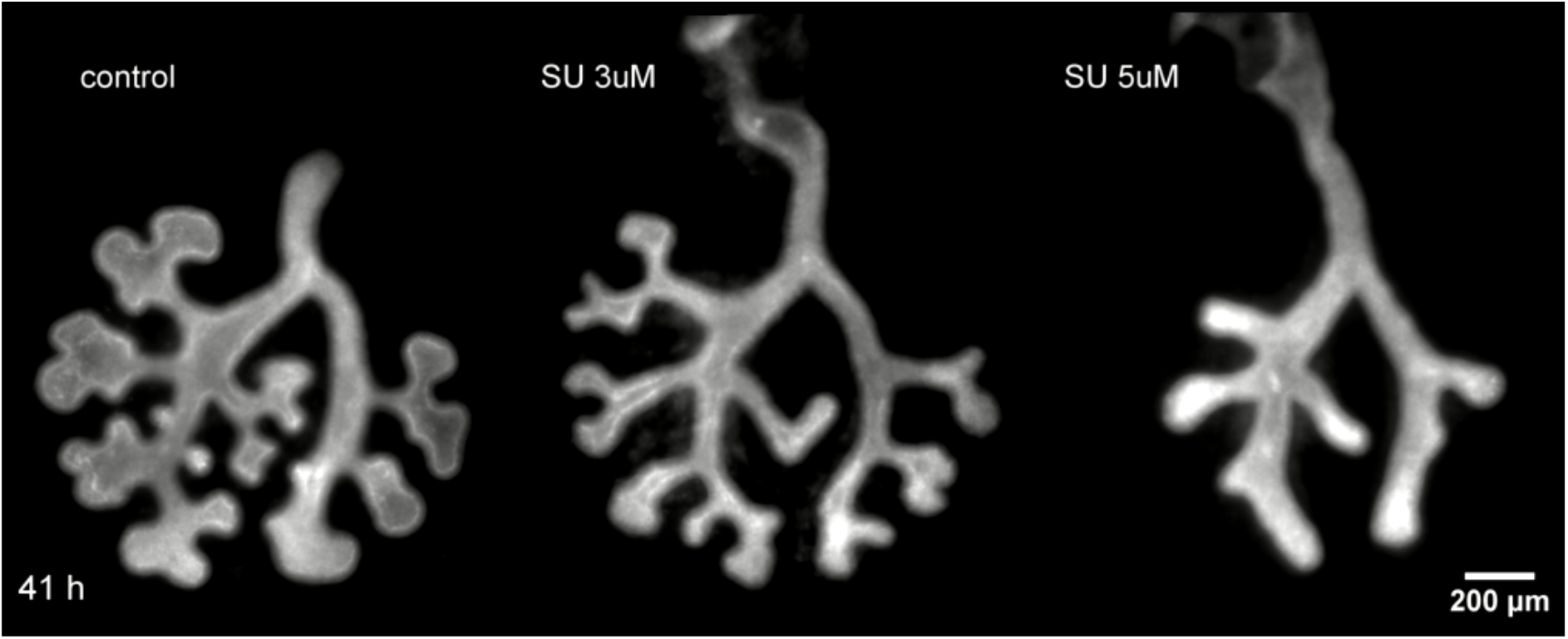
Lung explant cultures with FGFR inhibitor SU5402. Culture of embryonic lungs (E11.5) under control conditions and with the treatment of the FGFR inhibitor SU5402 at different concentrations for 48h. (AVI 8.16 MB).

**Supplementary Video 6:**
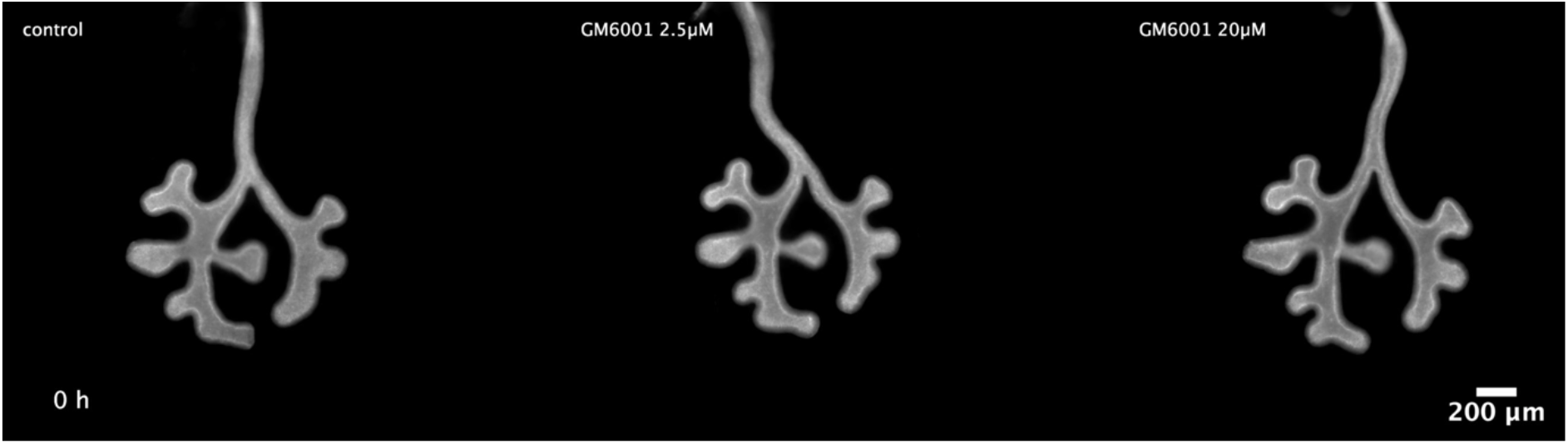
Lung explant cultures with MMP inhibitor GM60001. Culture of embryonic lungs (E11.5) under control conditions and with the treatment of the MMP inhibitor GM6001 at different concentrations for 60h. (AVI 5.3 MB).

**Supplementary Video 7:**
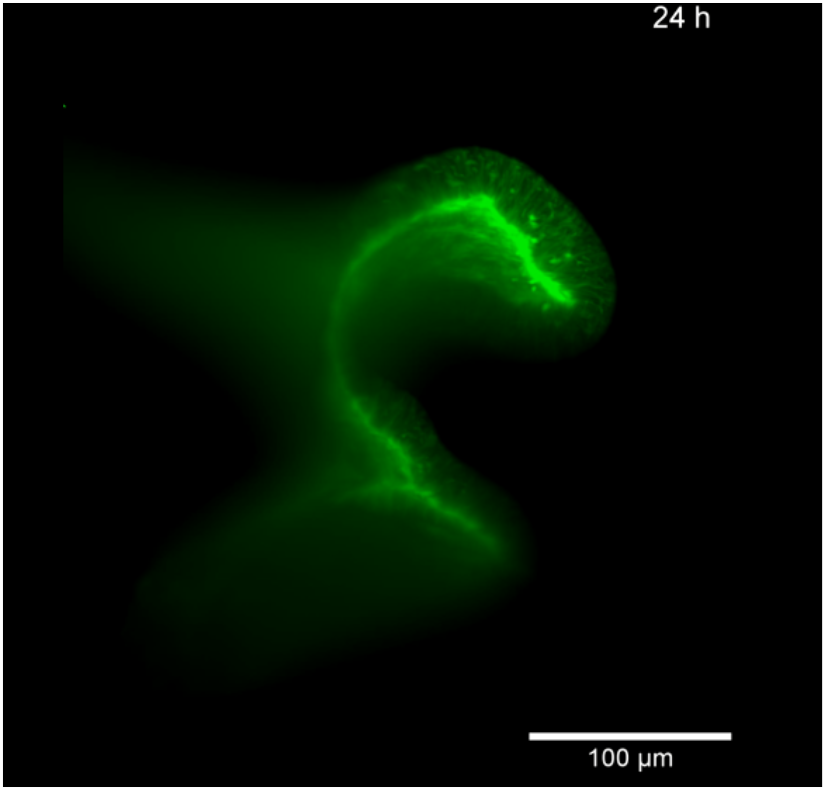
High-resolution light-sheet microscopy time-lapse imaging of lung bud elongation. Time-lapse movie showing the development of an E11.5 left lung rudiment carrying the Shh^GC/+;^ ROSA^mT/mG^ construct. The specimen was mounted in a hollow cylinder made from low-melting-point agarose and filled with matrigel to replicate the native microenvironment and promote near-physiological growth. Imaging was done using the Zeiss Z.1 Lightsheet system for 34h. (AVI 2.38 MB).

**Supplementary Video 8:**
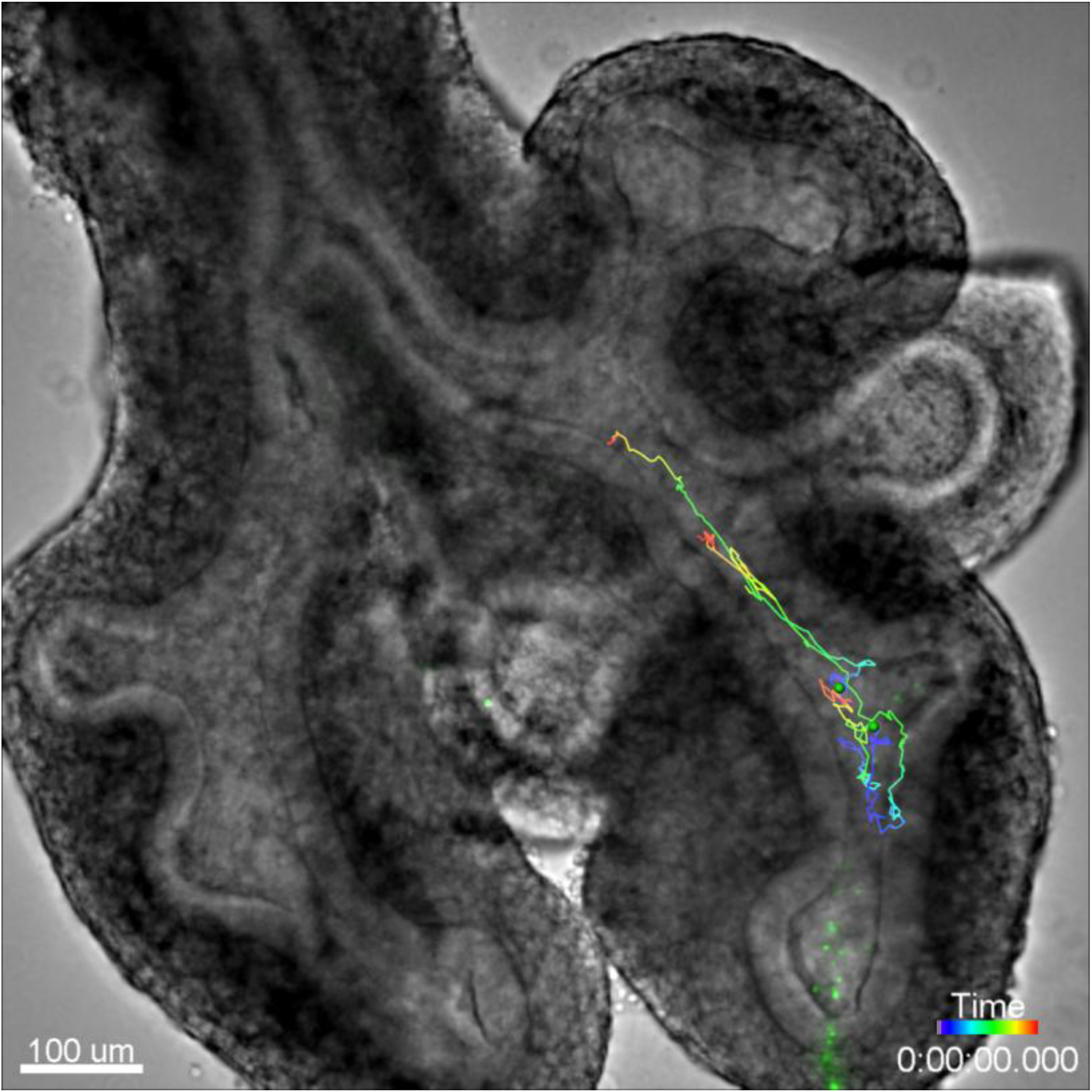
Tracking of beads in the lung lumen. Time-lapse movie of a 2 h spinning disk confocal acquisition (1 min time steps between frames) of fluorescent beads injected into the lung lumen. Bead tracks were generated in Imaris, green dots mark the detected bead at any given time point. (AVI 9.2 MB).

**Supplementary Video 9:**
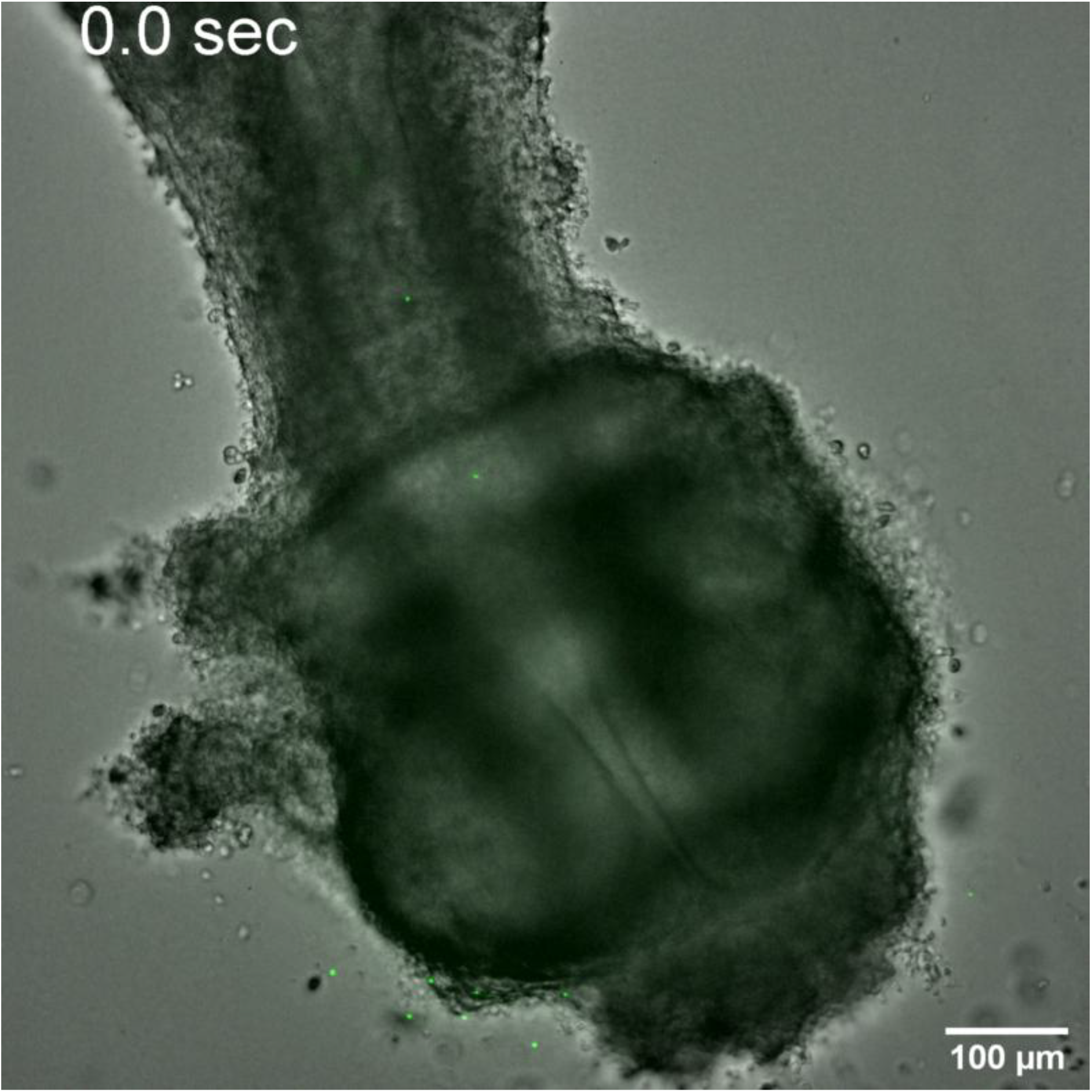
Outflow of fluid from the trachea opening. Spinning disk confocal imaging of fluorescent beads (488nm excitation channel, shown in green) overlayed with a brightfield image of the trachea of an E11.5 lung after 1h in low-volume culture. (AVI 2.5MB)

**Supplementary Video 10:**
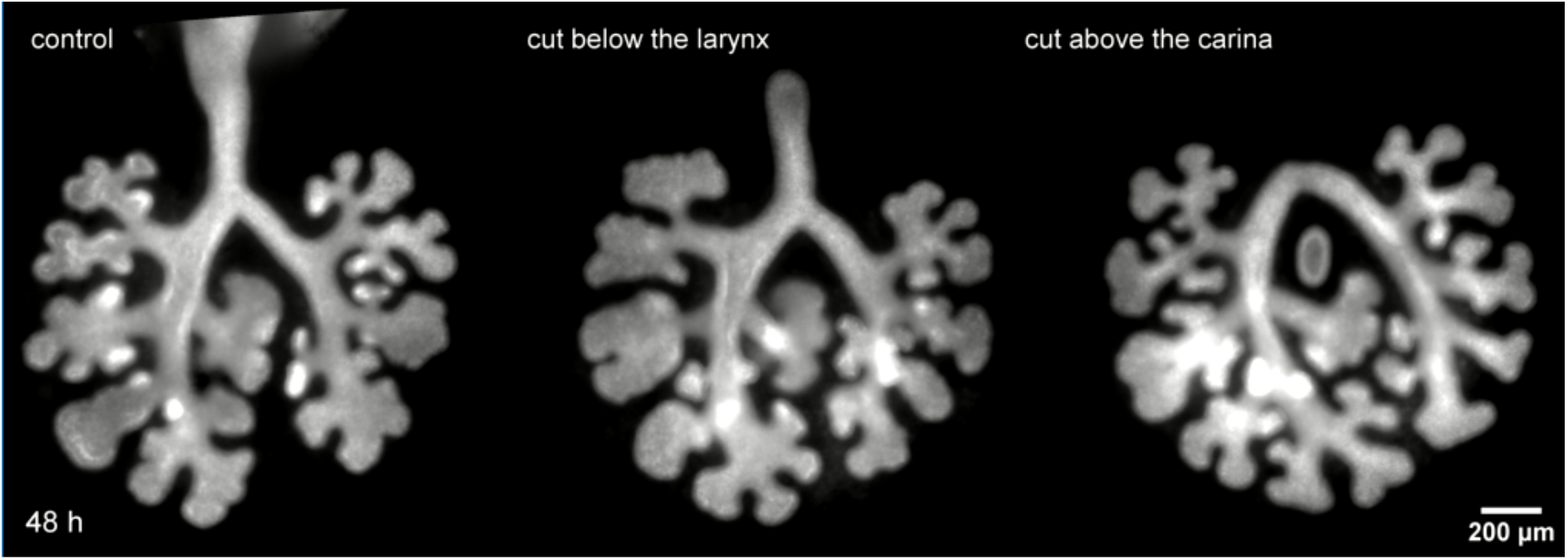
Lung explant cultures with intact and cut trachea. Culture of E11.5 embryonic lungs with intact trachea as control condition and with tracheas either cut below the larynx or above the carina for 48h. (AVI 11.8 MB).

**Supplementary Video 11:**
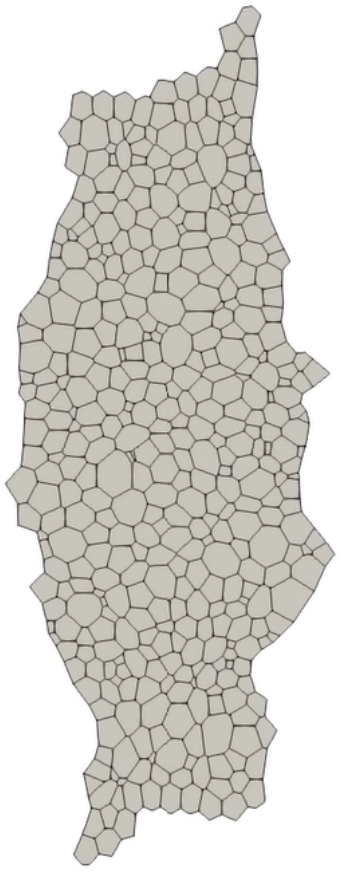
Tissue simulation with external force. Tissue growth simulation results of a vertex model when the initial configuration is subjected to a stretching force of 1.5 a.u. The parameters for the simulation are given in Supplementary Table 2.

